# NRP1 enhances endothelial barrier function via PDZ-mediated proximity to junction-related proteins

**DOI:** 10.64898/2026.11.27.769453

**Authors:** Tom P. Mitchell, Sammy El-Mansi, Vinothini Rajeeve, Claudio Raimondi, Thomas D. Nightingale

## Abstract

In the endothelium, neuropilin 1 (NRP1) is classically viewed as a co-receptor for vascular endothelial growth factor receptor 2 (VEGFR2), a pro-permeability, pro-angiogenic tyrosine kinase. During this response the PDZ-interacting domain of NRP1 is important for the maximal activation of VEGFR2 signalling. However, more recently NRP1 has been increasingly associated with impermeability responses and alternate roles.

We examined the effect of overexpression of wild type (WT) or a PDZ-interacting domain deletion mutant (ΔPDZ) of NRP1 on endothelial permeability. In the absence of vascular endothelial growth factor (VEGF)-A, NRP1 WT, but not ΔPDZ, caused a decrease in endothelial permeability. Consistent with a role in barrier function, knockdown of NRP1 increased endothelial permeability.

Using peroxidase proximity biotinylation coupled with mass spectrometry we identified proteins within a 20 nm labelling radius of either NRP1 WT or ΔPDZ in the whole cell, or in an enriched vesicular pool. This proteomic dataset was used to identify machinery associated with vascular permeability, but also provides insight to the other diverse aspects of NRP1 function.

Proteins significantly enriched at the cell surface with WT NRP1 include N-Cadherin, Junctional Adhesion Molecule-C and Integrin β3, all of which may disrupt junctional integrity. We propose a model whereby, in the absence of VEGF-A, NRP1 forms protein complexes which support the stability of endothelial junctions by sequestering these potentially destabilising factors. This is dependent on NRP1 PDZ-mediated interactions. However, further studies are required to identify the cytoplasmic machinery associated with these complexes and the impact of stimulation (by VEGF-A, shear forces and other stimuli) on vascular permeability.

**Highlights:**

- NRP1 enhances the *in vitro* barrier function of endothelial cells
- This role of NRP1 requires the presence of the PDZ-interacting domain
- Peroxidase proximity biotinylation allows identification of proteins differentially localised with WT or ΔPDZ NRP1
- NRP1 WT is proximal to junction-related proteins with the potential to disrupt established junctions
- We propose that in the absence of VEGF-A, NRP1 is present in a PDZ-dependent complex which prevents disruption of endothelial junctions

## Introduction

Neuropilin 1 (NRP1) was first identified as a neuronal marker (A5) in the optic system of *Xenopus laevis*^1^ and initially characterised as a cell adhesion protein^2^. However, this was superseded by independent discoveries of Kolodkin *et al*.^3^ and He and Tessier-Lavigne^4^ identifying NRP1 as the ligand binding component of class 3 semaphorin signalling. One year later NRP1 was shown to bind vascular endothelial growth factors (VEGFs) enhancing VEGF Receptor 1/2 (VEGFR1/2) signalling ^5^. This led to the view that NRP1 is predominantly a co-receptor enhancing the angiogenic potential of VEGFR2 signalling and the chemorepulsive effects of the class 3 semaphorins.

The development of murine knockout models seemed to confirm the NRP1/VEGFR2 interaction as NRP1’s biological role. Knockout models exhibited some strain-related differences (reviewed in Plein *et al*.^6^) however NRP1 knockout was universally lethal due to defective vascular development. Mechanistically, NRP1’s role in VEGFR2 signalling is mediated by the control of VEGFR2/NRP1 trafficking. NRP1’s C-terminal PDZ-interaction motif interacts with GIPC1 ^7^ and this interaction favours sorting into recycling rather than degradative pathways by protecting the complex from phosphatases ^8,9^. In agreement with this, deletion of the PDZ-containing cytoplasmic tail of NRP1 results in defective arteriogenesis ^10^.

However, this straightforward conclusion has been challenged on multiple fronts.

Firstly, it is increasingly evident that NRP1 is promiscuous as a co-receptor. NRP1 is also a co-receptor of c-Met^11^, transforming growth factor (TGF)-◻ receptors^12^, platelet-derived growth factor receptor ◻^13^, interleukin-13 receptor alpha1 ^14^, extracellular miRNAs ^15^ and the SARS-CoV-2 spike protein ^16^.

Secondly, NRP1 is required for non-angiogenic, and often ligand-independent functions. A non-exhaustive list includes: multiple roles in immunity and infection ^17–24^; stimulating local protein translation ^25^, regulating lipid uptake in multiple cells and promoting adipocyte differentiation ^26^; contributing to cancer metastasis ^27^, the production and regulation of the extracellular matrix ^28–30^; and promoting Hedgehog signalling ^31^ .

Thirdly, under the correct conditions NRP1 can inhibit angiogenesis and related processes. NRP1s potentiate class 3 semaphorin signalling in the endothelium ^32–38^. There is also evidence for a negative feedback loop - whereby VEGF-A stimulation leads to NRP1 cleavage and formation of anti-angiogenic cytoplasmic fragments ^39^. Recent evidence suggests that perivascular NRP1 acting in *trans* with endothelial VEGFR2 modulates the permeability effects of VEGF-A stimulation in animal models^40^.

Even in the absence of stimulation by an established NRP1 ligand, NRP1 modulates vascular permeability. Bossebouef *et al*. demonstrated exposure to shear force causes NRP1 to interact and stabilise VE-cadherin and adherens junctions - conferring an atheroprotective phenotype (though vascular permeability was not directly measured) ^41^.

To better understand the myriad functions of NRP1 and the importance of the PDZ-interacting motif we utilised proximity proteomics. This approach identifies nearby neighbours of HRP tagged NRP1 in live cells by catalysing the biotinylation of nearby proteins (within 20nm). By combining this approach with a membrane impermeant peroxidase inhibitor, ascorbate^42^, we separate proximal proteins present at the cell surface and within trafficked pools. Our data provides insight into the proteins surrounding NRP1 at the cell surface and in vesicular pools and their impact on endothelial junctions.

## Material and Methods

### HUVEC culture

HUVECs, isolated and pooled from up to four donors, were purchased from Promocell (Heidelberg, Germany) and used for experiments between passages 3-6. For general culture and passaging, cells were grown on dishes coated with 1% pig gelatin and maintained in M199 medium with Earle’s modified salts (Gibco, Grand Island, NY, USA) supplemented with 20% fetal bovine serum (Biosera, Nuaille, France or Labtech, Heathfield, UK), 10 U/ml heparin (Sigma Aldrich) and 30 μg/ml endothelial cell growth supplement (Sigma Aldrich). This medium was also used for live-cell imaging studies. Otherwise, post-lentiviral transduction, cells were switched into endothelial growth cell medium (Promocell) - a VEGF-A free formulation.

HUVECs were incubated at 37°C with 5% CO2 in humid conditions, except during the incubation step of lentiviral transduction where cells were maintained at 10% CO2 in otherwise identical conditions in a dedicated incubator in a biosafety level 2 lab.

### Production of Lentiviral particles

Human embryonic kidney 293T cells (a gift from Dr. James Whiteford, Queen Mary University of London, UK or the Lenti-X 293T cell line, Takara Bio Saint-Germain-en-Laye, France) were cultured in DMEM with Glutamax (Gibco) with 10% fetal bovine serum (Gibco). When producing lentiviral particles, cells were incubated in a dedicated incubator in a biosafety level 2 lab at 37°C with 10% CO_2_ in humid conditions.

To transfect cells for viral production, a mixture of 11 µg of plasmid expressing lentiviral packaging genes (pGagPol), 6 µg plasmid expressing the envelope (pMDVSVG), and 23 µg of the pLNT-SFFV plasmid containing the gene of interest were mixed with 24 mM polyethylenimine in 2ml OptiMEM (Gibco)

The mixture was maintained at room temperature for 15 minutes to allow complexes to form and then added to a T175 flask seeded with sub-confluent 293T cells. Virus-containing supernatant was removed, stored at 4°C, and replaced with fresh media for a further 24 hours. The two virus-containing supernatants were pooled, and syringe filtered with a 0.22 µM pore polyethersulfone filter. Four times concentrated polyethylene glycol precipitation solution was prepared (36% polyethylene glycol (6,000 Da), 1.6 M NaCl, filter sterilised) and diluted in the virus containing supernatant. This was stored at 4°C for 90 minutes, with tubes inverted every 30 minutes, before centrifugation at 1500 x g for 1 hour at 4°C. After centrifugation the supernatant was aspirated from pelleted viral particles and the pellet resuspended in OptiMEM. The viral titer was quantified using Lenti-X GoStix Plus (Takara Bio), prior to aliquoting and storage at -80°C. pGagPol, pMDVSVG and empty pLNT SFFV plasmids were a gift from Dr James Whiteford.

### Streptavidin Permeability Assay

HUVECs were seeded sub-confluently on biotinylated fibronectin (Cytoskeleton Inc., Denver, CO, USA) and transduced *in situ* with lentiviruses encoding NRP1 WT eGFP or NRP1 ΔPDZ eGFP. These were then grown to confluence over several days to permit the formation of functional cell-cell junctions. As a positive control cells were stimulated with 400 ng/mL of angiopoietin-1 (Peprotech, London, UK) for 30 minutes prior to changing to Dulbecco’s Phosphate Buffered Saline (PBS) with magnesium and calcium (to maintain junctional integrity) containing streptavidin-647 at 2 µg/mL.

After 3 minutes of streptavidin labelling, cells were fixed with 4% PFA and stained for nuclei (DAPI), GFP, VE-Cadherin and then imaged via confocal microscopy.

### NRP1 Knockdown and FITC-Dextran permeability assay

HUVECs from pooled donors (Lonza, UK) were cultured on 0.5 % gelatin in Endothelial Cell Growth Medium-2 with supplements (Lonza). For siRNA transfection, HUVECs were transfected with SMARTpool siRNA (Dharmacon, USA) against NRP1 (sense sequences: 5’-CGAUAAAUGUGGCGAUACU-3’; 5’-GGACAGAGACUGCAAGUAU-3’; 5’-GUAUACGGUUGCAAGAUAA-3’; 5’AAGACUGGAUCACCAUAAA-3’) or Silencer negative control siRNA (Life Technology, UK) using Lipofectamine RNAiMAX (Thermo Fisher Scientific, UK).

24 hours after transfection with si-NRP1 or si-Control, 5x10^5^ cells were seeded on 24-well transwell inserts (porous size 0.4µm) for a further 48 hours. Then, cells were starved in Endothelial Cell Basal Medium-2 for 3 hours, incubated with 70 kDa FITC-dextran (0.5mg/ml) in the top transwell chamber for 15 minutes. Fluorescence in the lower transwell chamber was read at 520 nm.

### Immunofluorescence staining

For immunofluorescent analysis cells were seeded onto glass coverslips sterilised with 70% ethanol and coated with 1% gelatin. Cells were washed with PBS, fixed with PBS 4% paraformaldehyde (PFA, supplied as methanol-free paraformaldehyde, Polysciences, Warrington, PA, USA) for 15 minutes and washed three times with PBS. Between fixation and staining, coverslips were stored at 4°C.

To stain cells, coverslips were rinsed in PBS 0.1% Triton X-100 (Sigma Aldrich), permeabilised with PBS 0.5% Triton X-100 for 10 minutes, then washed three times with PBS 0.1% Triton X-100. Blocking was performed with PBS 2% Bovine Serum Albumin (BSA, Sigma-Aldrich) 0.1% Triton X-100. Antibodies were diluted in PBS 2% BSA 0.1% Triton X-100 and primary and secondary incubation steps were for 30 minutes with 3 wash steps after each incubation as before. Coverslips were mounted on slides using ProLong Gold Antifade Mountant with DAPI (Thermo Fisher Scientific,). Images were captured using a Zeiss LSM 800 confocal microscope (Zeiss, Jena, Germany) and processed using the Fiji distribution of ImageJ ^43^.

### Western Blotting and protein pulldowns

Samples for western blotting were lysed on ice in radioimmunoprecipitation assay buffer (Sigma Aldrich or Merck Millipore, Burlington, MA, USA) and protease inhibitor cocktail (Sigma Aldrich). For proximity biotinylation experiments this was supplemented with 10 mM sodium azide to inhibit peroxidase activity. Proteins were separated by SDS-PAGE, then transferred to Immobilon-FL polyvinylidene fluoride membranes (Merck-Millipore) prior to antibody staining. Loading controls were multiplexed with proteins of interest by combining infra-red and HRP-conjugated secondaries on the same blot.

Co-immunoprecipitation of VEGFR2 with NRP1 WT eGFP was performed using GFP Trap beads (ChromoTek, Planegg-Martinsried, Germany). Beads were washed three times with lysis buffer tubes containing the beads and lysate were loaded onto a rotator overnight at 4°C. The next day beads were washed three times with PBS supplemented with protease inhibitor cocktail.

Biotinylated content was affinity purified with Pierce Streptavidin Agarose, or Pierce High Capacity Streptavidin Agarose (Thermo Fisher Scientific), depending on the scale of the assay. Pulldowns were performed as per GFP Trap (above) except that post-incubation washes were performed with PBS with 10 mM sodium azide (for western blot) or 25 mM ammonium bicarbonate buffer (for mass spectrometry). For western blotting, pulled down content was released from the beads by boiling with SDS-PAGE sample buffer.

### Peroxidase Proximity Biotinylation Assay

HUVECs were transduced with either equal titres of HRP-tagged constructs or empty vector controls as described above. For vesicular proximity biotinylation, 100 nM bafilomycin (Life Technologies) was added to the media for 4 hours prior to biotinylation to prevent lysosomal or autophagic degradation of vesicles.

Cells were fed 500 μM biotin tyramide (Iris Biotech, Marktredwitz, Germany), for 30 min at 37°C. The cells were then exposed to media supplemented with 1 mM H_2_O_2_ (Sigma Aldrich) in the presence (vesicular biotinylation) or absence (total biotinylation) of 50 mM freshly prepared ascorbate (Sigma Aldrich) for 1 minute. The biotinylation reaction was stopped by exchanging the media for 10 mM sodium azide, 10 mM ascorbate, 5 mM Trolox in PBS. Cells were then fixed or lysed, as described above.

### Mass spectrometry

Proteomics experiments were performed using mass spectrometry as reported ^44,45^. In brief, Immunoprecipated (IP) protein complex beads were digested into peptides using trypsin and peptides were desalted using C18+carbon top tips (Glygen Corporation, TT2MC18.96) and eluted with 70% acetonitrile (ACN) with 0.1% formic acid.

Dried peptides were dissolved in 0.1% TFA and analysed by nanoflow ultimate 3000 RSL nano instrument, coupled to an on-line Q Exactive plus mass spectrometer (Thermo Fisher Scientific). Gradient elution was from 3% to 35% buffer B in 120 min at a flow rate 250nL/min with buffer A being used to balance the mobile phase (buffer A was 0.1% formic acid in water and B was 0.1% formic acid in ACN). The mass spectrometer was controlled by Xcalibur software (version 4.0) and operated in the positive mode. The spray voltage was 1.95 kV and the capillary temperature was set to 255 °C. The Q-Exactive plus was operated in data dependent mode with one survey MS scan followed by 15 MS/MS scans. The full scans were acquired in the mass analyser at 375- 1500m/z with the resolution of 70, 000, and the MS/MS scans were obtained with a resolution of 17, 500. The mass spectrometry proteomics data have been deposited to the ProteomeXchange Consortium via the PRIDE ^46^ partner repository with the dataset identifier PXD087719 and 10.6019/PXD087719.

MS raw files were converted into Mascot Generic Format using Mascot Distiller (version 2.6) and searched against the SwissProt database (release 2020) restricted to human entries using the Mascot search daemon (version 2.5.0) with a FDR of ∼1% and restricted to the human entries. Allowed mass windows were 10 ppm and 25 mmu for parent and fragment mass to charge values, respectively. Variable modifications included in searches were oxidation of methionine, pyro-glu (N-term) and phosphorylation of serine, threonine and tyrosine. The mascot result (DAT) files were extracted into excel files for further normalisation and statistical analysis.

### Mander’s Co-localisation Coefficient Analysis

A single z plane was isolated from confocal z-stacks and individual cells were manually extracted, subjected to a Gaussian blur filter (σ=2.0) and then an appropriate automated thresholding method applied. A square image capturing the majority of the cell was generated from the blurred, thresholded images and split into individual channels. The subject channels were analysed with the Coloc2 plugin ^47^, then reanalysed with one channel rotated out of its true orientation to generate control values.

### Antibodies

See Supplemental Table 1

### Experimental Design and Statistical Rationale

For proteomic studies, the lysates of two 15 cm dishes per condition were pooled into a single technical replicate. 1.5-1.8 mg of lysate per technical replicate was loaded onto streptavidin beads.

Two technical replicates of each condition were analysed per biological replicate. Both technical replicates of each condition were run twice.

Mass spectrometry procedures were performed as described above, and normalisation processes were performed in Excel.

Enrichment of proteins was calculated as fold change between NRP1 WT or ΔPDZ HRP and empty vector control.

Student’s t-tests were performed comparing the average fold change between each NRP1 construct and control. These are shown in Supplemental Figure 2, however given the large number of comparisons, an alternative statistical analysis with a false discovery rate (FDR) correction was also employed.

Any protein showing an enrichment in the control rather than the NRP1 construct (ie. Fold Change <1) was removed as this is not due to specific biotinylation by NRP1-conjugated HRP.

Fold change data was then exported into GraphPad Prism 10 and examined for normality and lognormality. The data being lognormal, it was analysed using multiple lognormal t-tests adjusted using the two-stage Benjamini, Krieger, & Yekutieli FDR procedure method with a false discovery rate cut-off of 1% to determine significance.

Significant proteins were then analysed using the STRING database ^48^, applying an MCL clustering factor of 3, to help identify larger protein complexes or functional groups within the proxome.

For comparison between NRP1 WT or ΔPDZ proxomes (Figure 8 and Supplemental Figures 3-5-4) each run of the technical replicate was normalised to its equivalent empty vector control to create individual fold change values. This generates the same mean value as averaging the data first but allows reporting of variation (as mean ± standard error of mean). This is then shown on a log2 scaled axis to create a linear presentation b. As this data had already been normalised and statistically compared to the empty vector control, direct statistical comparison of the two proxomes was not considered appropriate.

## Results

### Generation of NRP1 overexpression constructs

To express tagged versions of NRP1 for both imaging and proteomics we identified an optimal site for tag insertion that did not interfere with localisation or function. We utilised an approach that was previously successful for tagging JAM-C ^42,49^ whereby we place the tag (eGFP or HRP) immediately before the transmembrane domain. This minimised disruption to both the extracellular and intracellular domain continuity (Figure 1A). This is particularly important in NRP1 as the N-terminal CUB domain of NRP1 regulates ligand binding, whilst the PDZ-interacting domain is C-terminal ^50^, which rules out common insertion sites at either the N or C terminus.

**Figure 1:**
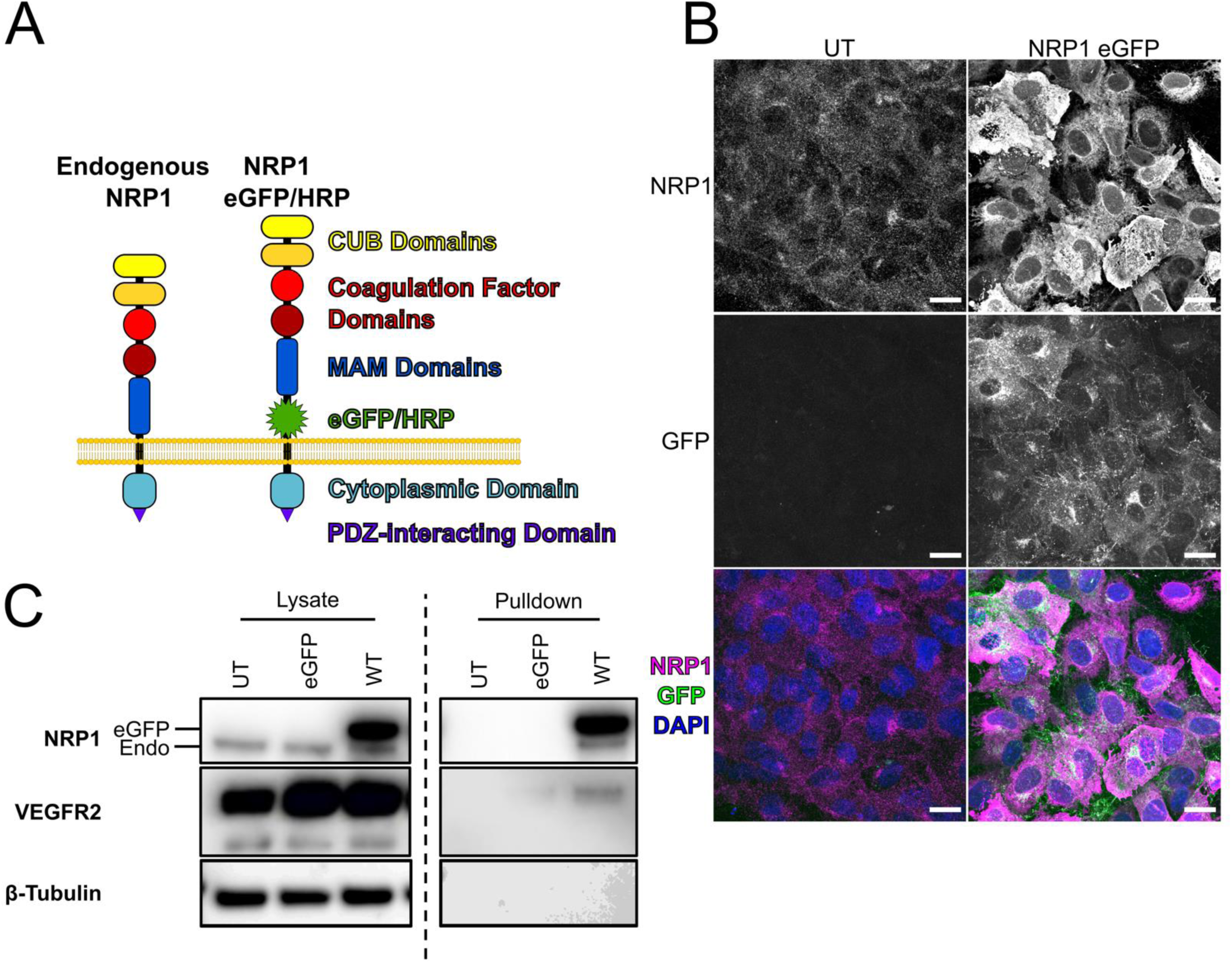
Insertion of juxtamembrane tags does not disrupt NRP1 trafficking. (A) Schematic of the structure of NRP1, with key functional domains and the insertion point of eGFP/HRP and PDZ interacting domains highlighted. (B) HUVECs were transduced with lentiviruses encoding NRP1 WT eGFP (or left untransduced, UT at a low titre. After growing to confluence, cells were fixed and stained for nuclei (DAPI), NRP1 and GFP then imaged via confocal microscopy. Scale bars 20 µm. (C) HUVECs were transduced as above, then lysed and GFP-tagged content (and interacting proteins) pulled down via GFP-Trap beads. Samples were western blotted and probed for NRP1, VEGFR2 and ꞵ-tubulin.

Following lentiviral transduction, we verified that NRP1 WT eGFP intracellular trafficking and intermolecular interactions were not disrupted. NRP1 eGFP transduced cells were reactive to an antibody targeting the NRP1 intracellular domain. eGFP signal localisation overlapped with antibody staining in the cell body but had more pronounced junctional localisation (Figure 1B).

GFP pulldown of NRP WT eGFP showed co-immunoprecipitation of both VEGFR2, and endogenous NRP1 - demonstrating successful co-receptor complex formation and homo-dimerisation of the eGFP-tagged construct (Figure 1C) ^51^.

### Overexpression of WT NRP1 construct decreases baseline endothelial permeability through its PDZ interacting domain

The capacity of NRP1 to promote VEGF-A-induced permeability has previously been established^52,53^. As most endothelial culture mediums contain VEGF-A to promote endothelial survival and identity maintenance ^54^, to avoid confounding effects of medium-derived VEGF-A, we performed permeability experiments in VEGF-A free formulations ^55^.

We overexpressed NRP1 WT eGFP and NRP1 ΔPDZ eGFP in HUVECs seeded on biotinylated fibronectin and performed a streptavidin permeability assay ^56^. This assay takes advantage of the high affinity interaction between streptavidin and biotin and the impermeability of streptavidin to the cell to assess the barrier function of a cellular monolayer. As a positive control for reducing vascular permeability we used preincubation with angiopoietin 1 ^57^.

We demonstrated that overexpression of NRP1 WT eGFP reduced vascular permeability to a level comparable to angiopoietin-1 stimulation (Figure 2A; 71.69% decrease in streptavidin positive area, *p*=0.0019). The effect was mediated through NRP1/PDZ protein interactions, as over-expression of NRP1 ΔPDZ eGFP to a similar level (Figure 2A; 6.00% increase in streptavidin positive area, *p*=0.9929) had no effect on vascular permeability.

**Figure 2:**
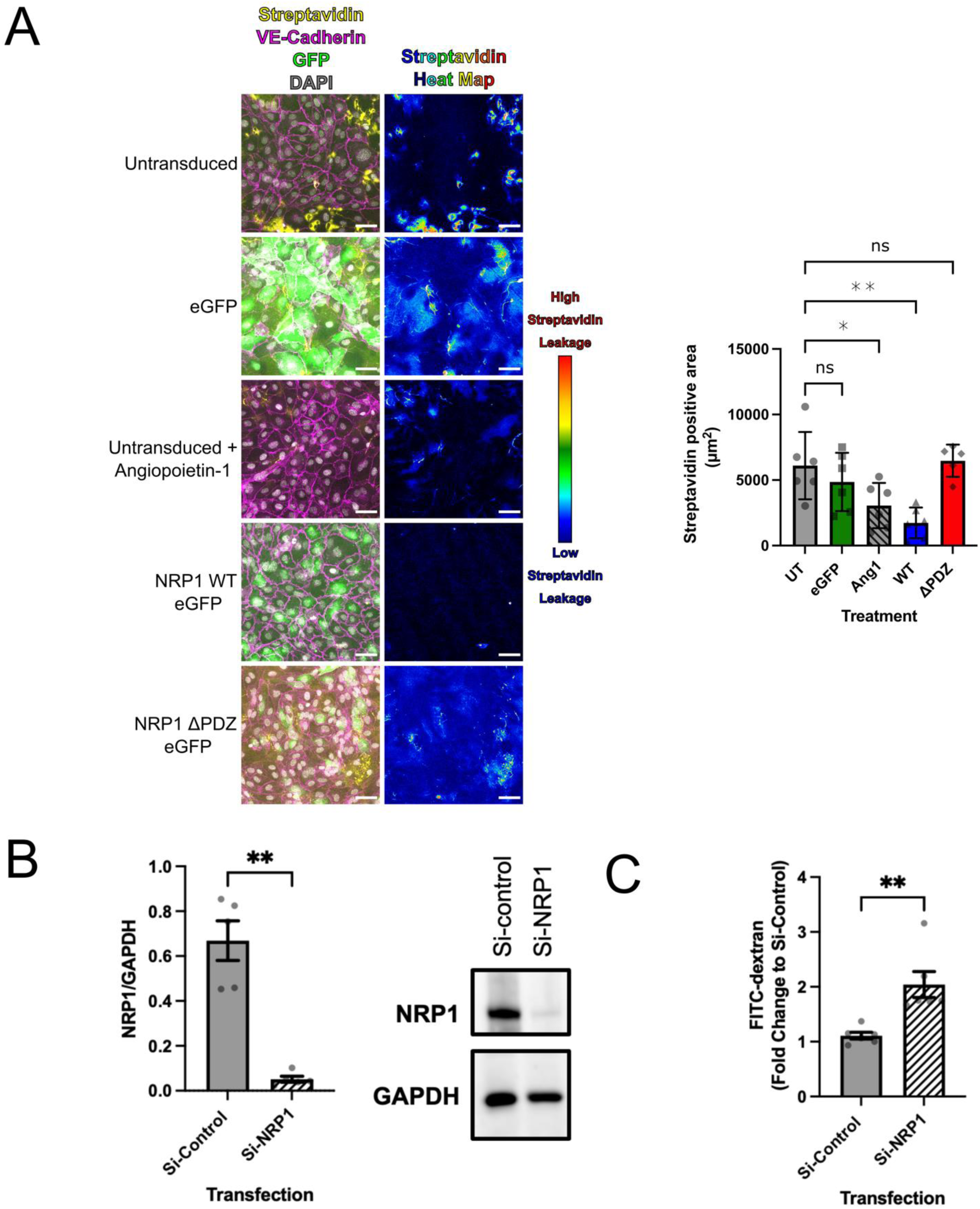
NRP1 positively regulates endothelial barrier function in VEGF-A free environments through its PDZ-interacting domain. (A) HUVECs were seeded on biotinylated fibronectin and transduced with lentiviruses encoding NRP1 WT eGFP or NRP1 ΔPDZ eGFP at an equivalent titre. After growing to confluence, cells were pre-incubated with 400 ng/mL of angiopoietin-1 for 30 minutes (Ang1) or left unstimulated. The culture media was then changed to PBS (with cations to prevent junctional disassembly) containing streptavidin-647 for 3 minutes, then fixed with 4% PFA. Cells were stained for nuclei (DAPI), GFP, VE-Cadherin and then imaged via confocal microscopy. Scale bars 20 µm. The streptavidin-positive area was quantified and averaged from 3 fields of views from 5-6 biological replicates per condition (some values excluded as outliers after ROUT analysis). The effect of each transduction or angiopoietin-1 pre-incubation was assessed via one-way ANOVA (*p =* 0.0010) with *post-hoc* Dunnett’s test. (B) HUVECs were transfected with control or anti-NRP1 siRNA and seeded on transwell inserts. *N* = 6 biological replicates, and protein reduction assessed by paired Student’s t-test. (C) Cells were switched to basal medium for 3 hours and 0.5mg/mL 70 kDa FITC-Dextran was added to the top chamber of the transwell and after 15 mins media was from the bottom well was collected and 520nm fluorescence measured. *N* = 6 biological replicates, and differences in fluorescence assessed by ratio paired t-test. Data shown as mean ± SEM

This change in permeability was not caused by gross changes in junctional morphology, VE-Cadherin levels or by changes in cell number (Supplemental Figure 1).

We further confirmed NRP1’s regulation of endothelial permeability using a knockdown study in an alternative permeability assay - monitoring FITC-labelled dextran passage through a transwell coated with a HUVEC monolayer.

SiRNA reduced HUVEC NRP1 content on average by 90.91% (Figure 2B), which led to a 1.84-fold increase in FITC-dextran leakage (Figure 2C; *p=*0.0160).

### Peroxidase proximity biotinylation reveals potential novel interactions, trafficking pathways and subcellular localisations of NRP1

To determine the difference in protein-protein interactions between WT and ΔPDZ NRP1 and the likely mechanism for the regulation of vascular permeability we utilised peroxidase proximity biotinylation. Compared to empty vector transduced cells (control) expression of HRP-tagged WT or ΔPDZ NRP1 resulted in a marked increase in biotinylation after 1 minute exposure to H_2_O_2_ (Figure 3A, “Total”, increase in streptavidin densitometry: WT NRP1 HRP - 92.65%; ΔPDZ NRP1 HRP - 79.12%). Cell surface biotinylation was blocked by the inclusion of cell-impermeant ascorbate, reducing a large amount of WT and ΔPDZ NRP1 HRP-mediated biotinylation (Figure 3A, “Vesicular”, decrease in streptavidin densitometry: WT NRP1 HRP - 38.59%; ΔPDZ NRP1 HRP - 35.44%). Immunofluorescent staining revealed that ascorbate-free (total) biotinylation assays produced a spider web-like network of labelling across the cell area (Figure 3B).

**Figure 3:**
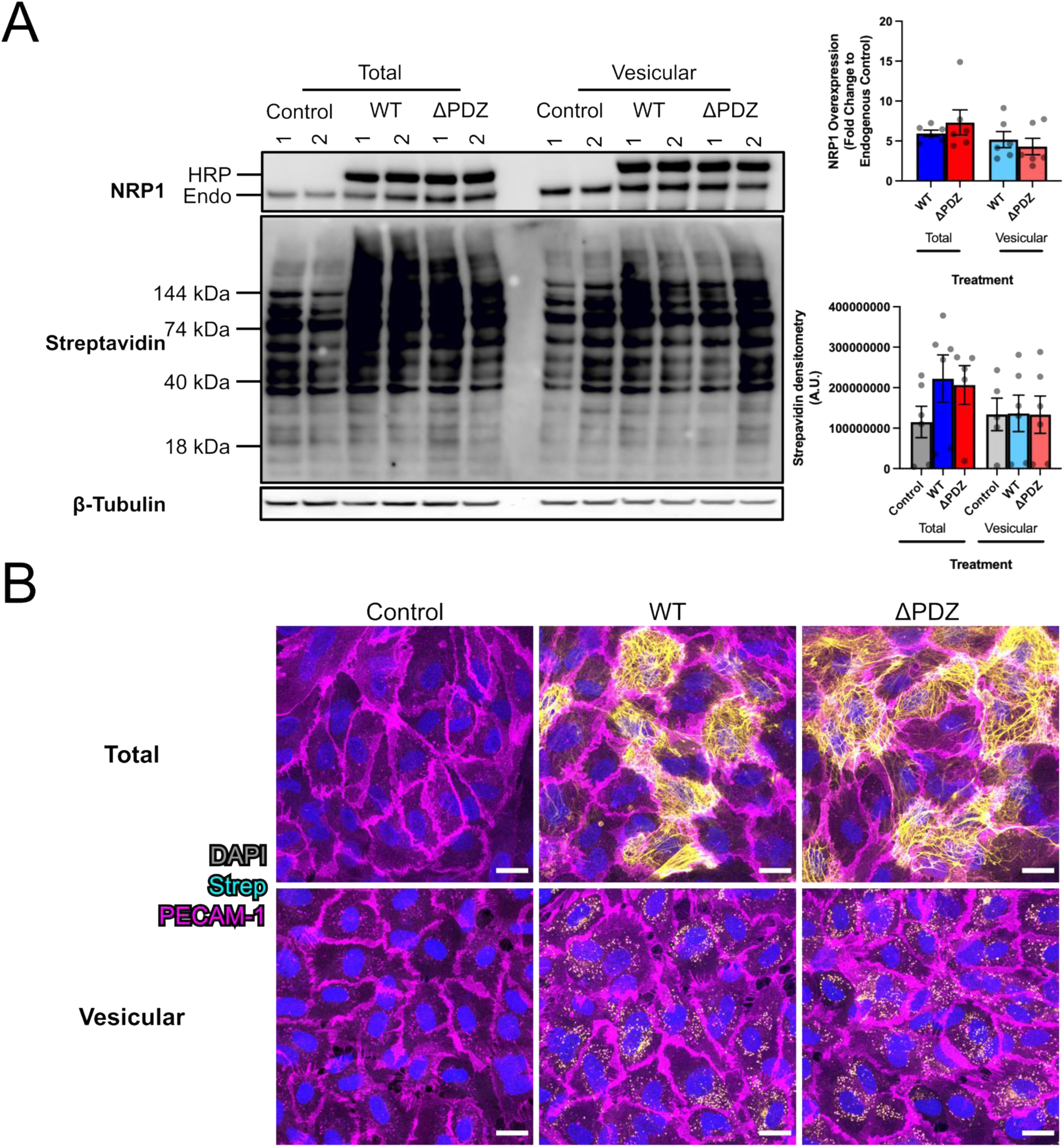
Validation of HRP proximity biotinylation via NRP1 HRP in both total and vesicular pools. HUVECs were transduced with lentiviruses encoding empty vector (Control), NRP1 WT HRP or NRP1 ΔPDZ HRP. Each group was split into 2 x 15 cm plates and seeded onto 2 x glass coverslips. After growing to confluence 2 x 15 cm plates and 1 x glass coverslip per group were incubated with bafilomycin A1 for four hours to increase vesicular content. All groups underwent proximity biotinylation - 30 minute incubation with biotin tyramide, followed by 1 minute stimulation with hydrogen peroxide to catalyse the labelling of proteins within a 20 nm radius with biotin. Bafilomycin A1 treated samples were stimulated in the presence of ascorbate to block cell surface labelling (vesicular), as opposed to untreated cells which were able to perform biotinylation at the cell surface (total). This process was carried out for two concurrent technical replicates per biological replicates (A) The 15 cm plates were lysed, and the 2 x 15 cm plates per technical replicate were pooled and western blotted for NRP1, streptavidin and ꞵ-tubulin (left). Right: Quantification of NRP1 HRP overexpression and biotinylation (via streptavidin densitometry) Data shown as mean ± SEM (B) To confirm the efficacy of bafilomycin A1 treatment and ascorbate incubation, glass coverslips were fixed and stained for nuclei (DAPI), biotinylated content (Strep), and PECAM-1 then imaged via confocal microscopy. Total biotinylation samples (top) exhibit a mesh-like network of surface biotinylation, whereas vesicular biotinylation (bottom) exhibit biotinylation only in endosomal-like punctae. Empty vector transduced samples exhibit no visible biotinylation. Scale bars 20 µm.

As previously established in our study of JAM-C ^42^, 4 hours of pre-incubation with the lysosomal/autophagosomal inhibitor bafilomycin increases the number of trafficking vesicles and labelling of co-trafficked proteins for mass spectrometry. This is particularly relevant for NRP1, which in some studies has been reported to have a very short half life (below 1 hour), primarily through lysosomal degradation ^58,59^.

Imaging of bafilomycin and ascorbate treated samples confirmed the ablation of surface labelling and enrichment instead of numerous endosome-like punctae (Figure 3B).

We scaled up the assay to allow sufficient protein for mass spectrometry and noted significant enrichment of proteins in all four conditions compared to the empty vector control (Supplemental Figure 2A). We excluded proteins where the empty vector control was enriched above the relevant NRP1 group (Supplemental Figure 2A/B). NRP1 specific proteins were then subjected to statistical analysis (incorporating FDR correction) to find significantly enriched proteins (Supplemental Figure 2B).

Significant hits were subdivided into ‘unique’ and ‘shared’ proteins along the various axes of comparison (Supplemental Figure 2C). For example, of the 101 significant hits in the NRP1 WT HRP total biotinylation dataset, 63 of these were also significant in the NRP1 ΔPDZ HRP total biotinylation dataset - resulting in 63 ‘shared’ hits with and leaving 38 ‘unique’ hits.

Of this same 101 significant hits in the NRP1 WT HRP total biotinylation dataset, 2 were also significant in NRP1 WT HRP total biotinylation dataset resulting in 2 ‘shared’ hits with and leaving 99 ‘unique’ hits.

In this study, we have primarily focused on ‘unique’ hits, as we are attempting to elucidate the mechanistic determinants for the change in vascular permeability. The data and comparisons are included (Supplementary Figures 3-5) and represent an important resource for other researchers in this field.

Comparatively few proteins increased their association with NRP1 following deletion of the PDZ-interacting domain. This is unsurprising given that PDZ-mediated interactions are key molecular cues, particularly at the cell membrane ^60^.

The total biotinylation analysis, (which is likely to reflect the proteins required for cell surface interactions, Figure 4) identified 38 proteins uniquely enriched within NRP1 WT HRP and only 6 uniquely enriched with NRP1 ΔPDZ HRP.

**Figure 4:**
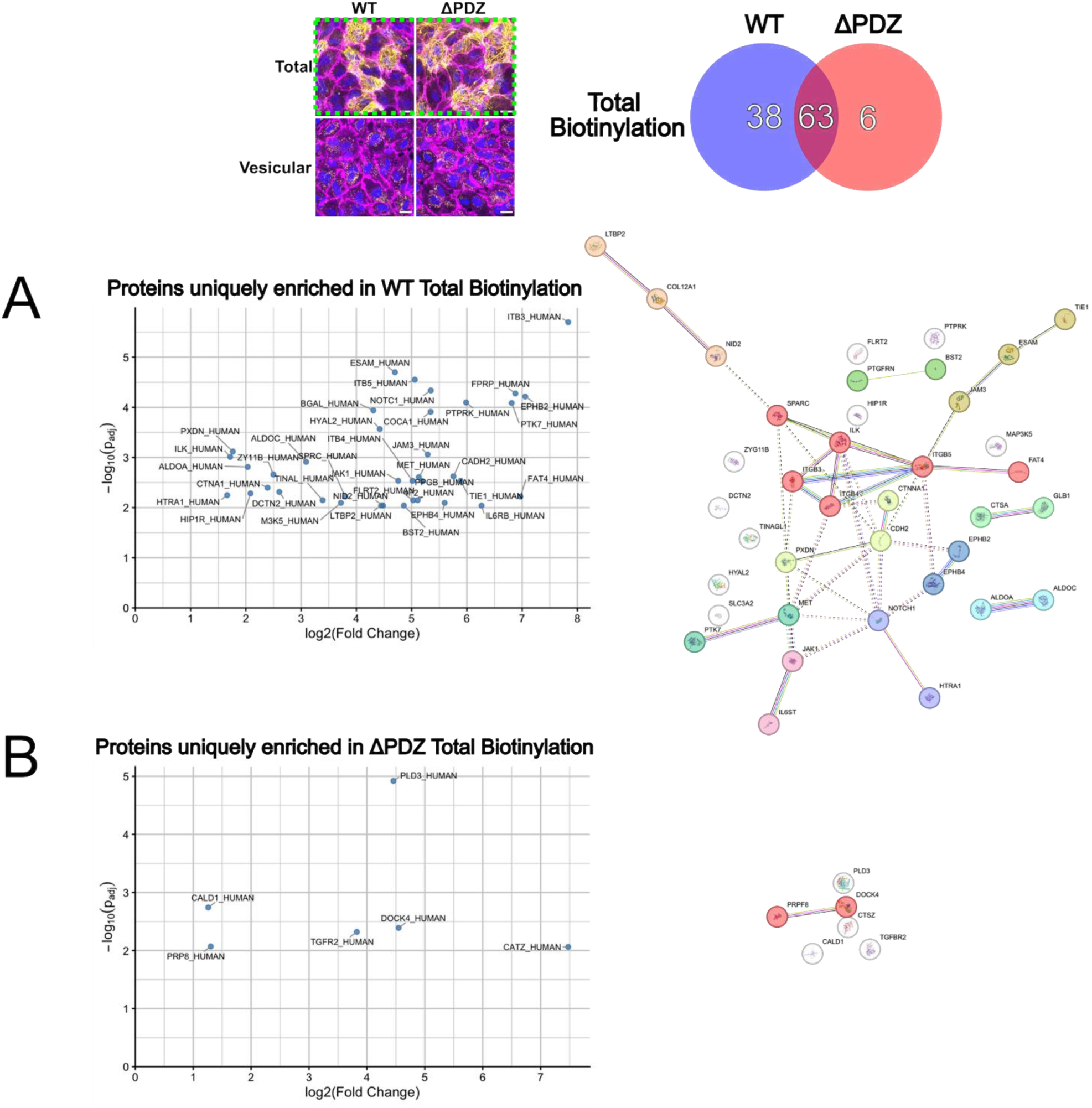
Proteins differentially enriched between NRP1 WT and ΔPDZ HRP in total biotinylation proteomic datasets. HRP proximity biotinylation and mass spectrometry was performed as described above. Filtered volcano plots are shown of proteins uniquely significantly enriched in total biotinylation proxomes of NRP1 WT (A) or ΔPDZ HRP (B) (fold change and *p*-values relative to empty vector control). STRING analyses of the enriched proteins were performed using an MCL clustering factor of 3. Underlying data is in Supplemental Table 3. Proteins enriched in both are found in Supplemental Figure 3.

WT-enriched proteins included clusters of cell adhesion and junction-related proteins. They also included proteins/clusters involved in NRP1 related functions such as extracellular matrix deposition and re-modelling ^28,29^ (such as nidogen 2 (*NID2*), collagen alpha-1(XII) chain (*COL12A1*), hyaluronidase-2 (*HYAL2*)), regulation of glycolysis (aldolase A (*ALDOA*) and C (*ALDOC*)) ^61–63^ and tyrosine kinase signalling (such as MET (*MET*), Janus Kinase 1 (*JAK1*), Tyrosine-protein kinase-like 7 (*PTK7*)) (Figure 4A) ^64,65^.

The only cluster identified by string analysis at the cell surface for NRP1 ΔPDZ HRP was between Dedicator Of Cytokinesis 4 (*DOCK4*) and Pre-MRNA Processing Factor 8 (*PRPF8*), this cluster is defined by co-expression and interaction of homologs in other species (Figure 4B). In agreement with published work TGFBR2 (*TGFBR2*) was identified as a proximal protein ^41^ , as were two lysosomally localised proteins, the 5’-3’ exonuclease PLD3 (*PLD3*) ^66^ and cathepsin Z (*CTSZ*) ^67^, supporting the idea that PDZ-mediated interactions may prevent NRP1 and co-trafficked cargo (such as TGFBR2) from entering lysosomal degradation pathways.

In samples pre-treated with bafilomycin and ascorbate, NRP1 WT HRP enriched proteins include a cluster of lysosomal proteins (lysosome-associated membrane glycoprotein 1 (*LAMP1*) and palmitoyl-protein thioesterase 1 (*PPT1*)) and clathrin associated proteins (AP-1 complex subunit mu-2 (*AP1M2*) and clathrin heavy chain 1 (*CLTC*)). This confirms that constitutive turnover of NRP-1 is via a clathrin dependent pathway ^68^ and that it is targeted for degradation in the lysosomes (Figure 5A).

**Figure 5:**
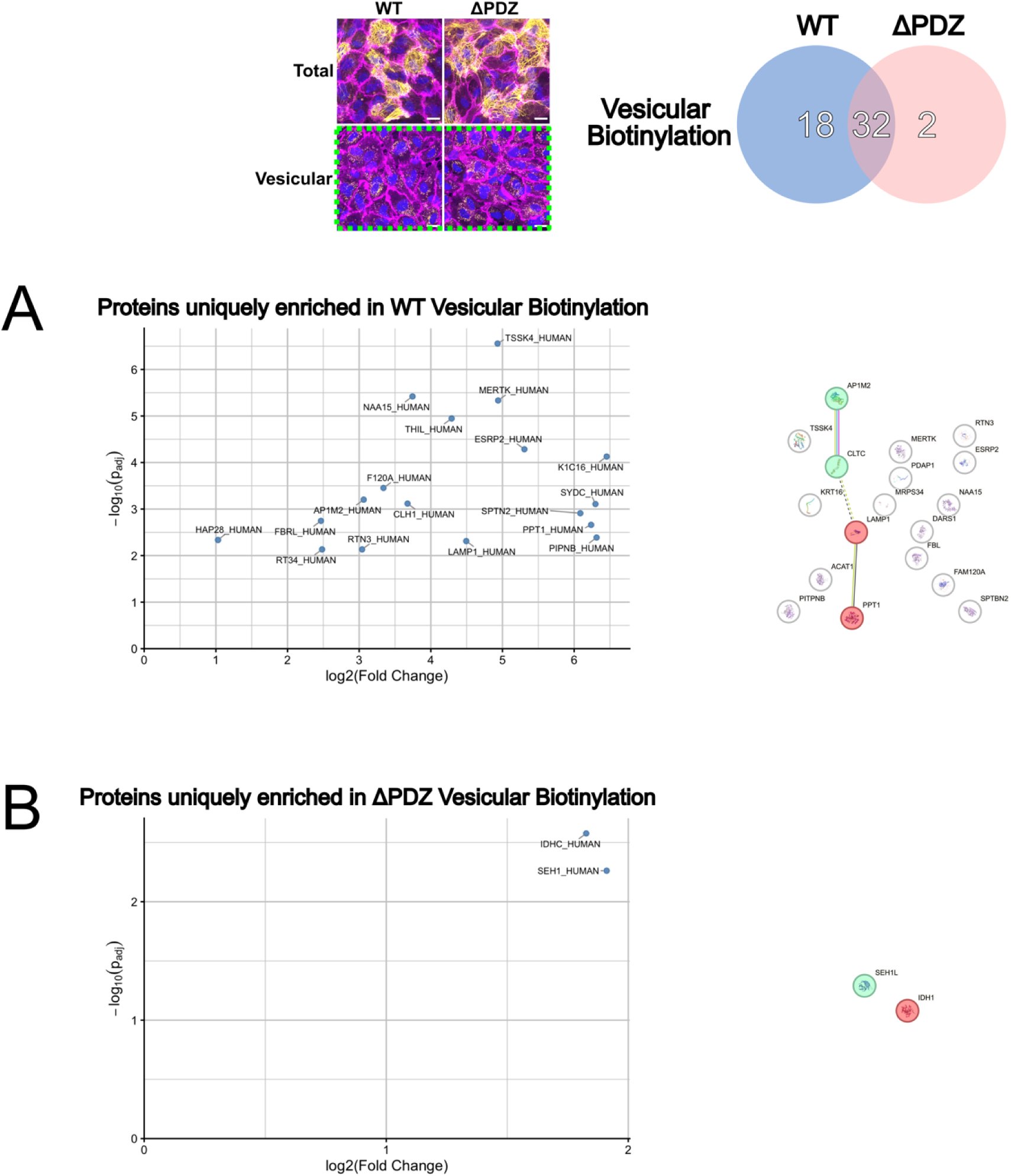
Proteins differentially enriched between NRP1 WT and ΔPDZ HRP in vesicular biotinylation proteomic datasets. HRP proximity biotinylation and mass spectrometry was performed as described above. Filtered volcano plots are shown of proteins uniquely significantly enriched in vesicular biotinylation proxomes of NRP1 WT (A) or ΔPDZ HRP (B) (fold change and *p*-values relative to empty vector control). STRING analyses of the enriched proteins were performed using an MCL clustering factor of 3. Underlying data is in Supplemental Table 4. Proteins enriched in both are found in Supplemental Figure 4.

Only two proteins were selectively enriched in the corresponding NRP1 ΔPDZ HRP samples - Isocitrate dehydrogenase [NADP] cytoplasmic (*IDH1*) and Nucleoporin SEH1(*SEH1L*) (Figure 5B). This highlights the importance of the PDZ-interacting domain for normal NRP1 localisation - and alternative subcellular localisations when these interactions are blocked. For example, as well as the cytoplasm, isocitrate dehydrogenase [NADP] cytoplasm is a peroxisomal protein ^69^. Nucleoporin SEH1 is a component of the nuclear pore complex ^70^ and NRP1 has previously been isolated from nuclear extracts of endothelial cells ^39^.

The majority of proteins uniquely enriched in total biotinylation of NRP1 WT HRP (compared to vesicular biotinylation) were junction-associated proteins, cell adhesion-related proteins and the semaphorin signalling pathway. However, there was also a cluster of five proteins associated with lysosomal storage disorders: N-acetylglucosamine-6-sulfatase (*GNS*; Sanfilippo syndrome type D ^71^ ); lysosomal alpha-mannosidase (*MAN2B1*; alpha-Mannosidosis ^72^ ); beta-galactosidase (*GLB1*; GM1 gangliosidosis ^73^ ) and both subunits of beta-hexosaminidase (*HEXA, HEXB*; two of the three GM2 gangliosidoses, Tay-Sachs disease ^74^ and Sandhoff disease ^75^) (Figure 6A).

**Figure 6:**
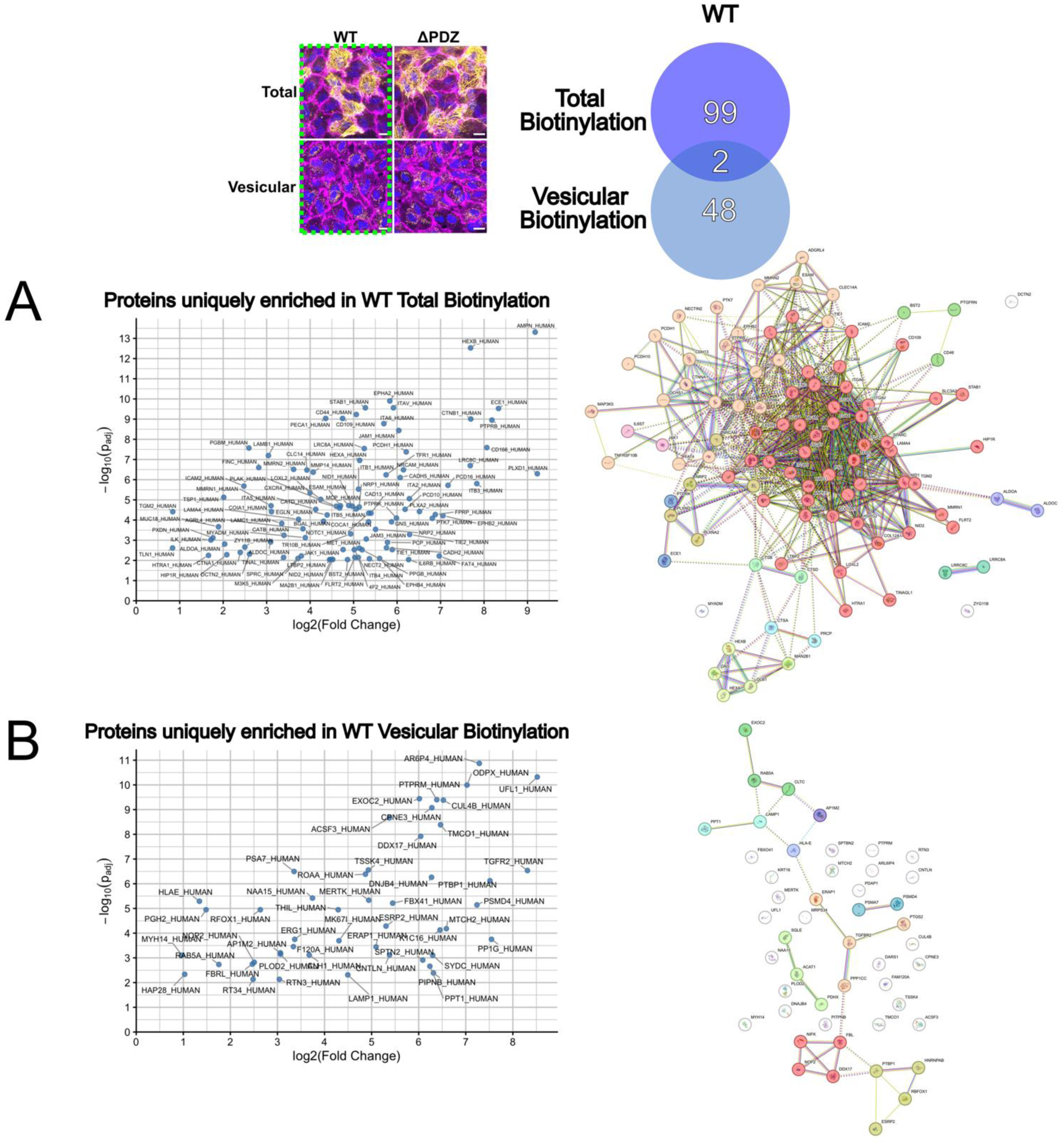
Proteins differentially enriched between NRP1 WT HRP total and vesicular biotinylation proteomic datasets. HRP proximity biotinylation and mass spectrometry was performed as described above. Filtered volcano plots are shown of proteins uniquely significantly enriched in NRP1 WT HRP total (A) or vesicular HRP (B) (fold change and *p*-values relative to empty vector control). STRING analyses of the enriched proteins were performed using an MCL

In the vesicular biotinylation data set NRP1 WT HRP exhibited clusters of proteins associated with an incredibly broad range of organelles (Figure 6B). As would be expected, these included components of the endolysosomal system (e.g. early endosomal marker Rab5a (*RAB5A*); lysosomal marker lysosome-associated membrane glycoprotein 1 (*LAMP1*)). As discussed above, AP-1 complex subunit mu-2 and clathrin heavy chain 1 are indicative of clathrin pit association but may also indicate transient association to the trans-Golgi network ^76^.

Less expected proximal proteins appear to be proteins from the mitochondria (Squalene monooxygenase (*SQLE*), Acetyl-CoA acetyltransferase, mitochondrial (*ACAT1*), Pyruvate dehydrogenase protein X component, mitochondrial (*PDHX*)); and the proteasome (proteasome subunit alpha type-7 (*PSMA7*); 26S proteasome non-ATPase regulatory subunit 4 (*PSMD4*)).

From a mechanistic perspective, STRING analysis identified a cluster of proteins functionally associated with the downregulation of TGF-β signalling (Figure 6B, orange cluster). There are also several proteins (red and olive clusters) associated with RNA processing - across the ribosomes, nucleus and cytoplasm. NRP1 has (as discussed above) previously been localised to the nucleus; binds to extracellular miRNAs ^15^; and regulates local translation in neurons through direct binding to mRNA and RNA-binding proteins (coupling them to ribosomes) ^25^.

However, for NRP1 ΔPDZ HRP only four of the five proteins identified in this cluster with WT were enriched. Instead, this cluster (olive cluster, Figure 7A) included lysosomal Pro-X carboxypeptidase (*PRCP*; also enriched with WT) and cathepsin Z (*CTSZ*; unique to ΔPDZ HRP). Both these proteins are peptidases that cut the C-terminal amino acid of proteins. The former regulates blood pressure by cleaving angiotensin II ^77^, and the latter is implicated in integrin regulation ^78^, angiogenic responses to VEGF-A ^79^ and pulmonary endothelial dysfunction leading to hypertension ^80^.

**Figure 7:**
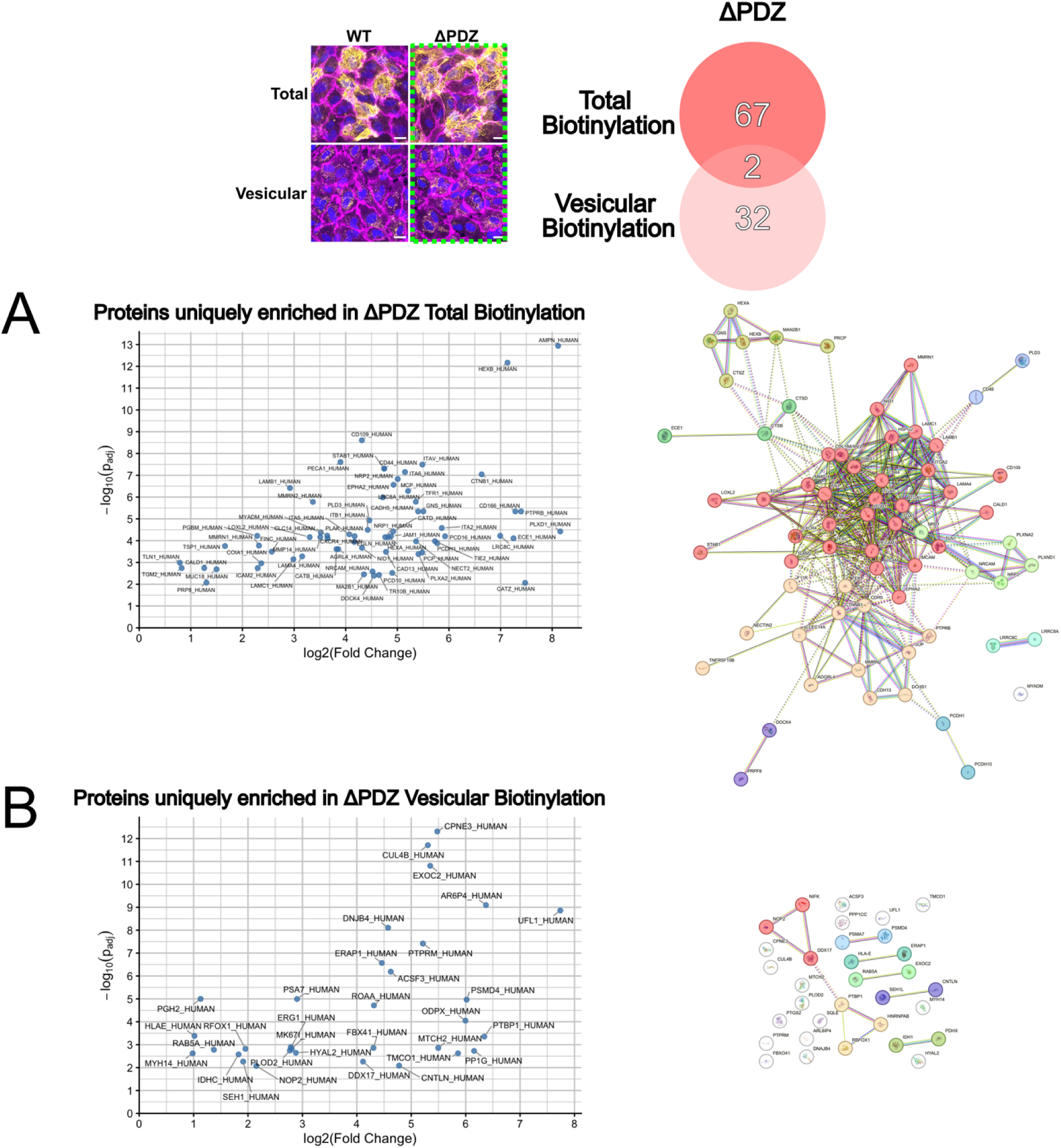
Proteins differentially enriched between NRP1 ΔPDZ HRP total and vesicular biotinylation proteomic datasets. HRP proximity biotinylation and mass spectrometry was performed as described above. Filtered volcano plots are shown of proteins uniquely significantly enriched in NRP1 ΔPDZ HRP total (A) or vesicular HRP (B) (fold change and *p*-values relative to empty vector control). STRING analyses of the enriched proteins were performed using an MCL

**Figure 8:**
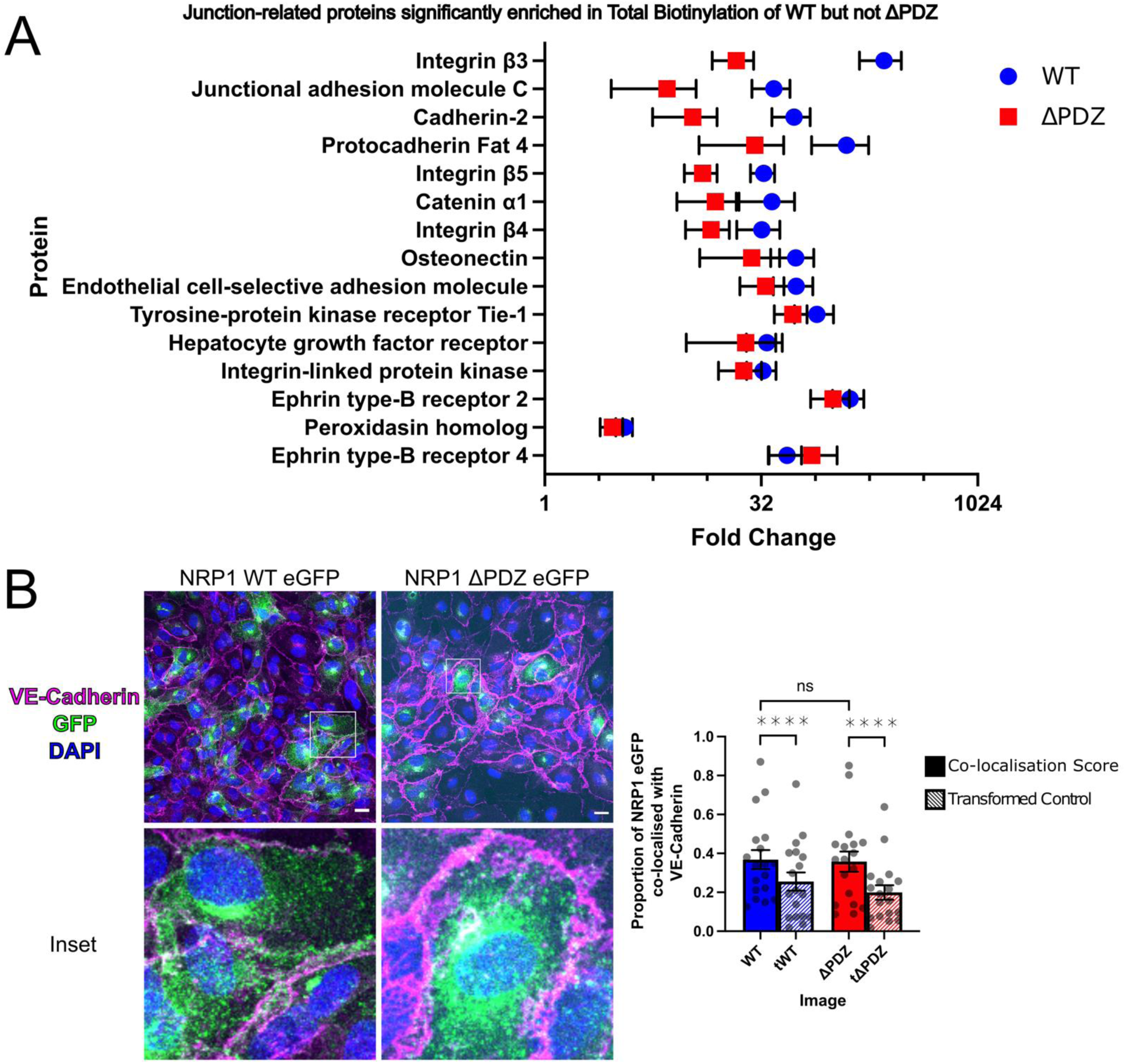
NRP1 WT HRP is significantly enriched with multiple endothelial junction and vascular permeability - related proteins compared to NRP1 ΔPDZ HRP, but both proteins are equally localised to the endothelial membrane. (A) Comparison of proteins which were significantly enriched in the NRP1 WT HRP total biotinylation dataset but not the NRP1 ΔPDZ HRP total biotinylation dataset (relative to empty vector control). Specifically, proteins identified in the red, dark goldenrod, green yellow, green and blue clusters of STRING analyses (Figure 6) and implicated in vascular permeability and/or cell contacts and adhesion (B) HUVECs were transduced with lentiviruses encoding NRP1 WT eGFP or NRP1 ΔPDZ eGFP at an equivalent titre. After growing to confluence, cells were fixed and stained for nuclei (DAPI), GFP and VE-Cadherin then imaged via confocal microscopy. Magnified images of the area outlined in white are included to demonstrate co-localisation of eGFP and anti-VE-Cadherin antibody. Manders co-localisation co-efficients of WT NRP1 eGFP and NRP1 ΔPDZ eGFP with VE-Cadherin were calculated for aligned images and transformed (t) controls. Both constructs were significantly enriched over transformed controls (paired Student’s t -test) but with no significant difference between constructs (unpaired Student’s t -test). *N*3 fields of views each from 6 biological replicates. Scale bars 20 µm.

The enrichment of two of the three subunits (leucine-rich repeat-containing protein 8A and C; *LRRC8A, LRRC8C*) which formulate a variant of the endothelial volume-regulatory anion channel specifically found to regulate blood pressure ^81^ are further suggestive of potential roles for NRP1 in endothelial control of vascular tone. However, leucine-rich repeat-containing protein 8A and C heteromers are also associated with cellular responses to viral infection ^82^, and this finding must be considered in the context of lentiviral transduction used to express the constructs.

As there are only 2 uniquely enriched proteins for vesicular biotinylation of NRP1 ΔPDZ HRP (relative to WT NRP1 HRP) most of the unique proteins and clusters relative to total biotinylation have already been discussed above. This likely reflects defective trafficking of the ΔPDZ NRP1. There are several proteins in this group, however, which may be indicative of post-translation modifications of either NRP1 itself or proximal proteins (Figure 7).

These included two proteins involved in phosphatase activity (serine/threonine-protein phosphatase PP1-gamma catalytic subunit, *PPP1CC*; receptor-type tyrosine-protein phosphatase mu, *PTPRM*); components of two distinct E3 ubiquitin ligase complexes (F-box only protein 41, *FBXO41*; and Cullin-4B, *CUL4B*) and an E3 ligase of ubiquitin fold modifier 1 (E3 UFM1-protein ligase 1, *UFL1*).

### The PDZ-interacting domain is responsible for the formation of junctional protein complexes around NRP1, but not its membrane localisation

Our approach identified clusters of cell adhesion and junction-related proteins, (Figure 4A, red, dark goldenrod, green, yellow, green and blue clusters) that were significantly enriched in NRP1 WT HRP but not NRP1 ΔPDZ HRP sample. These represent the likely mediators of the observed change in endothelial permeability.

Whilst these proteins were significantly enriched in NRP1 WT HRP but not NRP1 ΔPDZ HRP samples, in order to assess which proteins control the permeability effect we compared the relative fold change over control for these proteins in the total biotinylation datasets (Figure 8A)

The greatest differences in fold change (WT vs ΔPDZ, mean fold change ± standard error of the mean) were observed in integrin β3 (228.09 ± 73.91 vs.21.44 ± 6.85), JAM-C (39.15 ± 11.58 vs. 7.10 ± 4.20) and cadherin 2 (aka N-cadherin 54.06 ± 16.02 vs. 10.70 ± 5.07). A further 5 were accompanied by a shift of >2 fold towards NRP1 WT.

This change in proximal proteins was not due to a gross change in the junctional localisation of NRP1 following deletion of the PDZ-interacting domain. Mander’s co-localisation analysis ^83^ demonstrated that both NRP1 WT and ΔPDZ eGFP were significantly associated with the junctional marker VE-Cadherin; with no significant differences in co-localisation score between them (Figure 8B). As such, we hypothesise the changes in permeability and proximity are due to alterations in protein complexes formed within the plasma membrane itself.

## Discussion

Our data supports broader roles for NRP1 than acting as a co-receptor for VEGFR2 and other receptors, particularly in the context of permeability. When VEGF-A was absent from the culture medium, NRP1 strengthened the endothelial barrier through PDZ-dependent mechanisms.

### PDZ-dependent enhancement of endothelial barrier function by NRP1

Our proximity proteomics highlighted clusters of junctional and cell adhesion proteins that were significantly enriched with WT but not ΔPDZ NRP1 HRP. Although these were significant in total biotinylation samples they were not significantly enriched in the vesicular pool - these effects are therefore determined by interactions or proximity at the cell surface, rather than by altered trafficking. The three which exhibited the greatest difference were integrin β3, JAM-C and N-Cadherin.

All three of these proteins mirror the complexity of NRP1 - as well as being promoters of cell-cell or cell-matrix interactions; they also promote vascular permeability.

Integrin αvβ3 interacts with NRP1 in endothelial cells during angiogenesis - isolating it from interactions with VEGFR2, and potentially sequestering VEGF-A itself ^84^. However, a positive synergistic signalling relationship between integrin β3 and VEGFR2 in response to VEGF-A stimulation has also been observed ^85^. As such, the relevance of this interaction is unclear, particularly in our low VEGF-A model of vascular permeability.

In the specific context of vascular permeability, knockout or inhibition of β3 integrins is associated with reduced barrier function ^86–88^. Mechanistically, Su *et al,* demonstrated that treatment with the anti-permeability factor sphingosine-1 phosphate caused the translocation of αvβ3 to cortical (sub-plasma membrane) sites - where it engaged with extracellular matrix ligands to form focal adhesions and stabilised the actin cytoskeleton to increase barrier function ^86^.

This model was further refined by Alghisi *et al*. ^89^ who found that selectively inhibiting extracellular matrix interactions with αvβ3 led to its displacement from focal adhesions into the cell junctions. There it increased vascular permeability by displacing VE-Cadherin through recruitment of Src kinase and phosphorylation of VE-Cadherin ^89^.

Incorporating this model with our key observations (a PDZ-dependent decrease in vascular permeability in NRP1 overexpression; and the proximity of integrin β3 to WT NRP1 at the cell surface) suggests a model whereby PDZ-mediated interactions with NRP1 retain β3 in focal adhesions (especially cortical) preventing it from translocating to the junctions to destabilise them. This model assumes PDZ-proteins act as scaffolding intermediaries. A likely candidate for further investigation would be syntenin, which interacts with Src kinase and promotes the formation and activity of Src/focal adhesion kinase complexes ^90^.

Intriguingly, JAM-C has previously been shown to form a complex with integrin β3 in the regulation of vascular permeability ^91^. Li *et al*. ^91^ showed in unstimulated cells JAM-C and integrin β3 co-localise and immunoprecipitate and following pro-permeability stimuli JAM-C rapidly translocates to the junctions. Both overexpression of JAM-C and pro-permeability stimulation led to reduced junctional integrin β3, and reduced adhesion to integrin β3 ligands or VE-Cadherin. Although the molecular mechanism for this isn’t clear, a complex between JAM-C and the PDZ-protein afadin was found to negatively regulate the activity of Rap1b - a promoter of integrin activation ^91^.

Li *et al.’*s displacement of β3 from the junctions appears to contradict the observations of Alghisi *et al*. and there are other conflicting reports of the effects of changes in JAM-C and integrin β3 expression and localisation on permeability ^86,92^.

What is consistent in these reports is that JAM-C and integrin β3 can therefore modulate vascular permeability through cytoskeletal re-arrangements and modified VE-Cadherin function. These shared features and their established complex formation is suggestive of a shared mechanism, likely involving the inactivation and release of integrin β3 from cortical focal adhesions by JAM-C/afadin/Rap1b. The role of NRP1 in such a mechanism would be unclear. As NRP1’s direct interaction with integrin β3 has been established ^84^, and both JAM-C and NRP1 have PDZ-interacting domains it is feasible for it to form a complex with either or both.

As well as identifying JAM-C and integrin β3 as proximal to NRP1 in this study, we previously established via JAM-C HRP peroxidase proximity biotinylation that integrin β3 and NRP1 are proximal to JAM-C at the cell surface in endothelial cells ^42^. However, the scale of difference was markedly biased towards NRP1 (fold change over control = 4638.05), compared to integrin β3 (fold change over control = 4.08). NRP1 was also identified as sharing a trafficking pathway with JAM-C while integrin β3 did not - suggesting that the JAM-C/NRP1 interaction is more common JAM-C/integrin β3 interaction.

Combining this observation with the differing permeability effects observed in this and other studies, it seems likely that NRP1 helps stabilise JAM-C and integrin β3 in separate pools, opposing their potential pro-permeability effects (individually or as a complex). Given the PDZ-dependency of their proximity and the permeability effects in this study, PDZ-scaffolding proteins are key candidates for further investigation to elucidate this interaction. As discussed above, for integrin β3 syntenin would be a candidate ^90^; and for JAM-C: afadin ^91^, Par3 ^93^ and zonula occludens (ZO)-1 ^93^ and -2 ^94^.Identifying afadin or ZO-1 as NRP1 interactors would be particularly interesting, as these also have roles in coordinating N-Cadherin function ^95^.

N-Cadherin is a protein with an atypical role in endothelial permeability. In immature endothelial monolayers it acts as an endothelial junction protein, but is replaced by VE-Cadherin as the endothelium matures ^96^. In mature monolayers it preferentially strengthens heterotypic cell interactions (e.g. between endothelial and smooth muscle cells) ^97^. Overexpressing N-Cadherin leads competition with VE-Cadherin for p120 catenin - disrupting the VE-Cadherin junctions^96^.

As such, it appears likely that NRP1 would stabilise N-Cadherin in non-junctional locations to prevent disruption of VE-Cadherin junctions.

### Non-permeability roles for NRP1

Beyond permeability, and in keeping with the pleiotropic nature and diverse functions of NRP1 we identified a large number of proteins indicative of localisation from the nucleus to the mitochondria; and with implications for diseases as varied as hypertension and lysosomal storage disorders.

Whilst this study focused on unique hits, proteins which are enriched in multiple pools (Supplemental Figures 2-4) can also provide interesting mechanistic insights. For example, Bosseboeuf *et al*. identified that in endothelial cells under flow NRP1 sequesters transforming growth factor beta receptor 2 (TGFR2) into intracellular pools ^41^. In this study, TGFR2 is uniquely enriched with NRP1 ΔPDZ HRP compared to NRP1 WT HRP in both total (fold change over control = 14.18, *p_adj_* = 0.0048) and vesicular (fold change over control = 199.26, *p_adj_* = 0.0084) biotinylation datasets - suggesting that PDZ-mediated interactions are essential for NRP1’s regulation of TGFBR2.

We also identified clusters of proteins associated with lysosomal storage disorders in both NRP1 WT (5 proteins) and ΔPDZ (4 proteins) HRP total biotinylation samples. As this assay identifies proximal proteins not direct interactions, this may only indicate a shared lysosomal localisation - especially as NRP1 undergoes incredibly rapid degradation in lysosomes (with a half life below 1 hour in some studies) ^58,59,98^.

However, there are significant overlaps in NRP1 functions and pathological features of these disorders: neuronal function ^99^; processing of cell-surface and extra-cellular matrix; mitochondrial function and iron regulation ^100–102^.

Furthermore, a previously published RNASeq dataset identified significant reductions in both beta-hexominadase subunits (*HEXA:* 45.01% reduction, *p_adj_* = 0.0173; *HEXB:* 46.93% reduction, *p_adj_* = 0.0002) and beta-galactosidase (36.68% reduction, *p_adj_* = 0.0053) transcripts in endothelial cells where NRP1 was knocked down ^103^. Together this suggests an underappreciated role for NRP1 in neuronal metabolism which may have implications for disease.

In NRP1 WT HRP vesicular biotinylation samples a cluster of proteins associated with RNA processing was identified. One possibility is that these proteins are proximal within cytoplasmic ribonucleoprotein granules, such as P-bodies or stress granules.

Alternatively, given several nuclear localised proteins were identified in proximity with NRP1, this cluster could represent non-cytoplasmic ribonucleoprotein granules such as Cajal bodies or the nucleolus ^104^.

As well as proteins which may form functional complexes with NRP1 we identified proteins which may regulate NRP1 or its interacting proteins via post-translational modification. Of these serine/threonine-protein phosphatase PP1-gamma catalytic subunit; receptor-type tyrosine-protein phosphatase mu; F-box only protein 41; Cullin-4B and E3 UFM1-protein ligase 1 were proximal even after deletion of the PDZ-interacting domain.

There remains the possibility that NRP1 may undergo post-translational modifications via these proteins including phosphorylation^105,106^, ubiquitylation^107,108^ and UFMylation^109^.

### Conclusions, study limitations and future work

Overall, the PDZ-dependent permeability effect we observed is likely due to complex formation of NRP1 with PDZ proteins (key candidates being GIPC1 syntenin, afadin, ZO-1) to stabilise key junction related proteins - e.g. integrin β3, JAM-C and N-cadherin. These could stabilise junctions directly (integrin β3 being maintained in focal adhesions and promoting a favourable actomyosin state), or indirectly by preventing disruption of junctions (JAM-C’s effects on VE-Cadherin and integrin β3; N-Cadherin competing with VE-Cadherin) (Figure 9).

**Figure 9:**
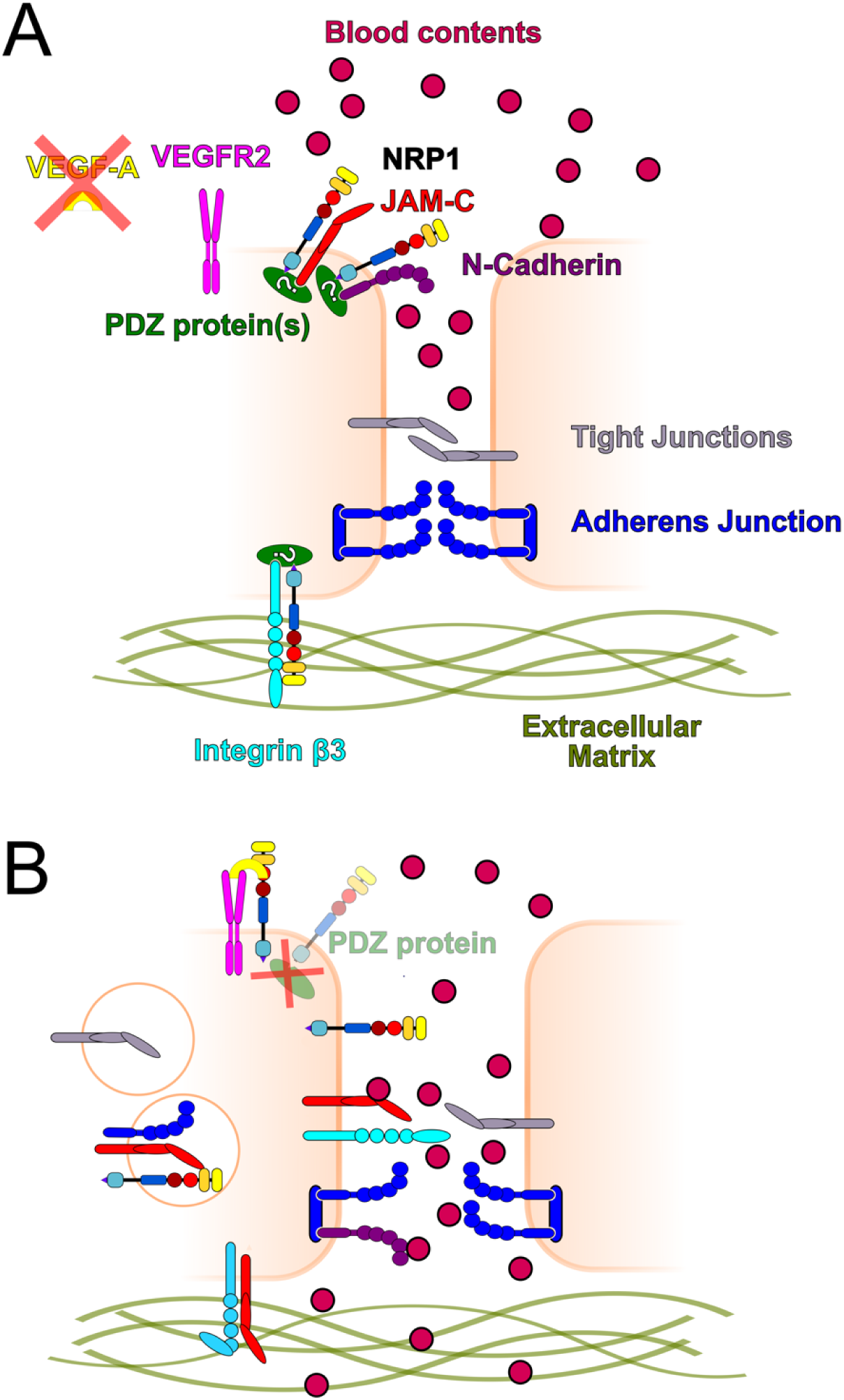
Proposed model of NRP1’s effect on vascular permeability. (A) In the absence VEGF-A, NRP1 associated with junction-related and and cell adhesion proteins. These interactions are dependent on its PDZ-interacting domain and therefore likely involve unknown PDZ proteins as scaffolds. The complexes stabilise and prevent disruption of the endothelial junctions, enhancing the barrier function and reducing *in vitro* permeability – which would translate into passage of blood contents from the vessel into the tissue. (B) In the presence of VEGF-A, or the absence of PDZ-mediated interactions NRP1’s these complexes disassemble. This leads changes in localisation (shown here for JAM-C and N-Cadherin); changes in activation (shown here for Integrin β3 and additional downstream effects (including on the cytoskeleton and VE-Cadherin). The net effect is an increase in vascular permeability.

This model suggests the stabilisation occurs in distinct subcellular compartments. Whilst we observed no significant changes in localisation or junctional morphology between our WT or mutant NRP1, there is a striking difference in the biotinylation pattern between NRP1 HRP and JAM-C HRP from our previous study ^42^. NRP1 HRP exhibits biotinylation in a spider web-like pattern across the entirety of the cell, whereas JAM-C HRP biotinylation is restricted to the junctions. Given that integrin β3, JAM-C and N-Cadherin all have important roles at the apical/basal cell surface (e.g. extracellular matrix binding ^89^, leukocyte adhesion ^49^, and smooth muscle cell interactions ^110^, respectively) it seems plausible that NRP1 may stabilise these proteins in their apical/basal localisation as opposed to their lateral/junctional localisation. However, this is technically challenging to evaluate by microscopy in HUVECs, which have a maximum height of ∼1.5 µm ^111^.

To verify protein complexes, both proximity identified proteins like integrin β3 and putative PDZ-interactors such as GIPC1, approaches such as immunoprecipitation or knockdown coupled to functional assays are required.

Unbiased identification of PDZ-scaffolding proteins, and/or other cytoplasmic components of protein complexes, could be achieved using an ascorbate peroxidase 2-based peroxidase proximity biotinylation approach. This alternative peroxidase retains full activity in the reducing environment of the cytoplasm, and we have previously utilised it to identify the cytoplasmic machinery associated with JAM-C ubiquitylation ^42^.

Finally, our experiments were performed in the absence of stimulation. In particular, our study and model utilises a VEGF-A free environment. The addition of VEGF-A seems likely to dissociate these proposed complexes, and the potential junctional destabilisation effects of proteins like N-Cadherin may contribute to VEGF-A’s pro-permeability effects.

To test this hypothesis, and the broader roles of NRP1 it would be beneficial to repeat this experimental approach in the presence of VEGF isoforms, semaphorins, shear etc. This would take advantage of the short labelling times required for peroxidase proximity biotinylation to examine changes in proximal proteins before and after stimulation ^112^.

Whilst this study is unable to fully elucidate the role of the NRP1 and its PDZ-interacting domain in the regulation of endothelial barrier function, we have established a permeability effect of NRP1 in the absence of VEGF-A, and used proximity labelling and proteomics to identify proteins potentially involved. The data represent a useful baseline for further studies on the regulation of vascular permeability and the broader study of NRP1; and the lentiviral constructs produced can be used to investigate other NRP1 functions in the endothelium and other cell types.

## Acknowledgments

The authors acknowledge the CMR Advanced Bio-Imaging Facility of Queen Mary University of London for the help and advice with microscopy. The authors thank their funders and collaborators.

This work was supported by Barts Charity Project Grant MGU05434 and by Queen Mary University of London.

S.E.-M was supported by British Heart Foundation Fellowship FS/IBSRF/26/25340

C.R was supported by British Heart Foundation Fellowship FS/16/22/32045 and Barts Charity Seed Grant G-002671.

## Authorship Contribution

T.P.M. and T.D.N. developed the methodology and wrote the original draft of the manuscript; T.P.M, S.E.-M, V.R., C.R. performed the investigation; T.D.N. supervised the study; and all authors reviewed and edited the manuscript.

## Declaration of generative AI and AI-assisted technologies in the manuscript preparation process

During the preparation of this work the authors used ChatGPT in order to generate scripts for the R package ggplot2. These were used solely to modify the appearance of plots generated by the package (e.g. changing font sizes, modifying axis width etc.), and never to modify the underlying data. All data manipulations, such as log changes, were performed in Excel and imported into the R environment for plotting. After using this tool, the authors reviewed and edited the content as needed and take full responsibility for the content of the published article.

## Abbreviations

ANOVA -: Analysis of Variance
FDR -: False Discovery Rate
HRP -: Horseradish Peroxidase
HUVEC -: Human Umbilical Vein Endothelial Cell
JAM -: Junctional Adhesion Molecule
NRP1 -: Neuropilin-1
PBS -: Phosphate Buffered Saline
PFA -: Paraformaldehyde
SEM -: Standard Error of the Mean
TGFR2 -: transforming growth factor beta receptor 2
UFM1-: ubiquitin fold modifier 1
VEGF -: Vascular Endothelial Growth Factor
VEGFR -: Vascular Endothelial Growth Factor Receptor
ZO -: Zonula Occludens

---

**Supplemental Figure 1:**
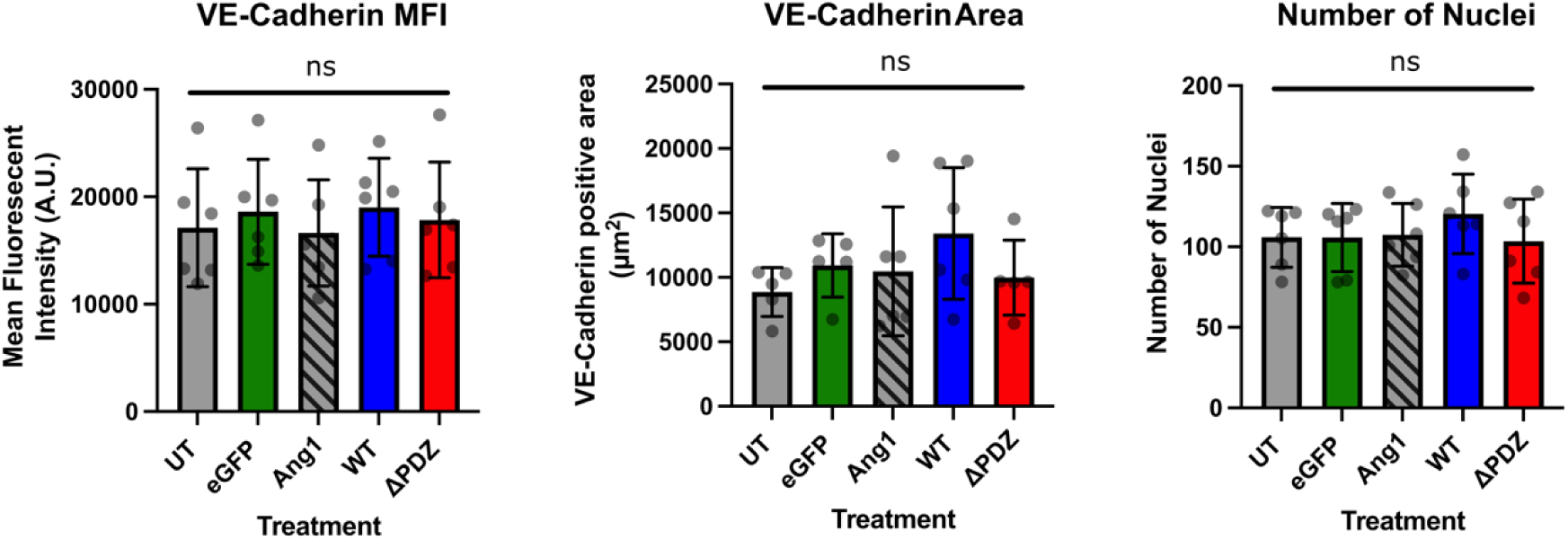
VE-Cadherin Quantification. Streptavidin permeability changes in Figure 2B are not due to changes in VE-Cadherin levels (assessed via MFI, *p =* 0.9185), junctional status (assessed by VE-Cadherin area, *p =* 0.3969) or cell number (assessed via nuclei, *p =* 0.6951) – all assessed by one way ANOVA. Data shown as mean ± SEM, *N* = 6.

**Supplemental Figure 2:**
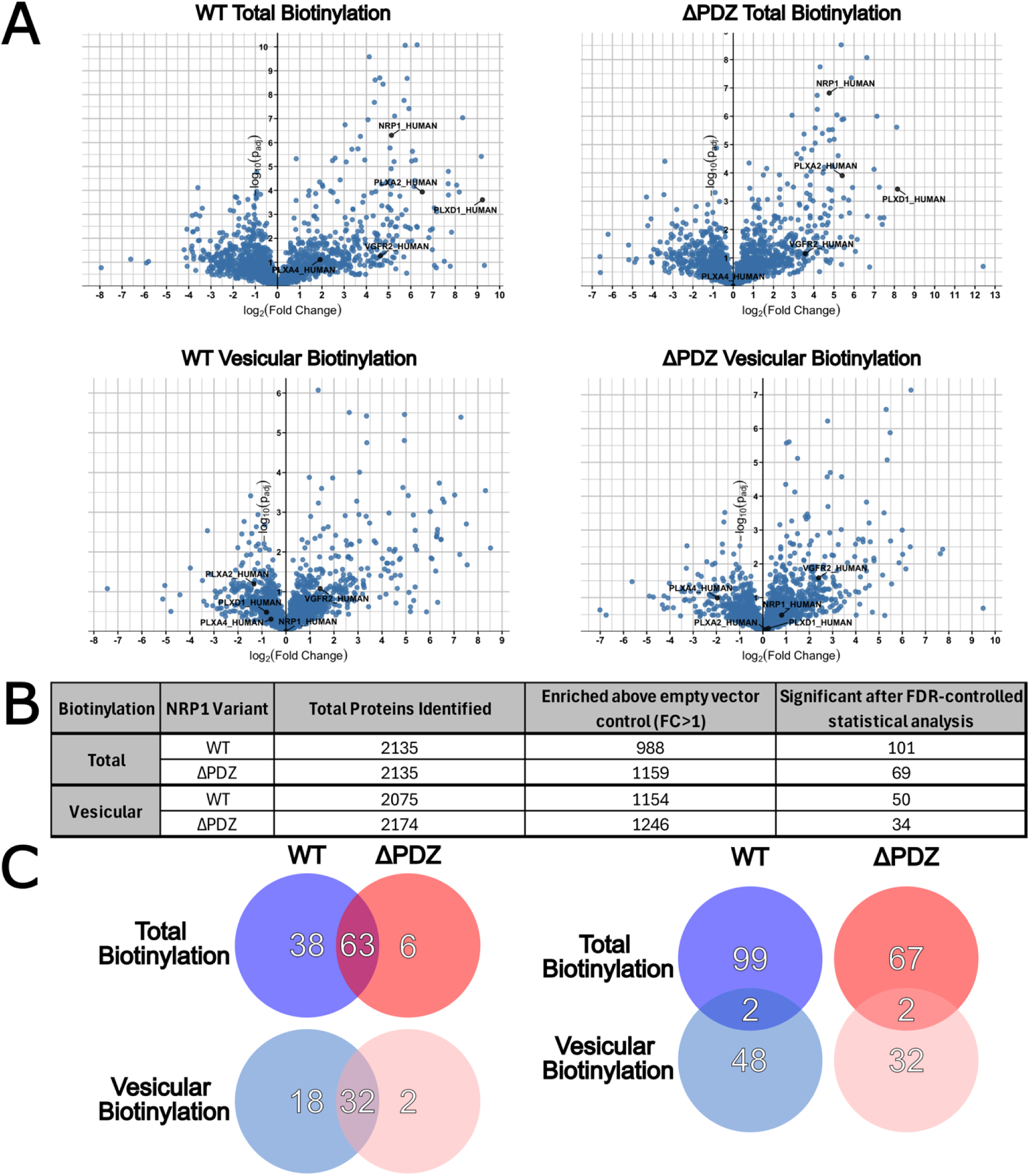
Mass spectrometry identifies a large number of proteins from NRP1 HRP proximity biotinylation experiments, necessitating contextual filtering and conservative statistical analysis. HRP proximity biotinylation was performed as described above. For mass spectrometry studies, 2 x 15 cm plates per technical replicate were pooled and after confirming equivalent expression of constructs and successful biotinylation via western blot, an equal amount of protein (1.5-1.8 mg) was loaded onto streptavidin-coated beads to pull down biotinylated content overnight at 4°C. Biotinylated proteins were identified via on-bead tryptic digest followed by mass spectrometry. Three biological replicates were analysed, with the mass spectrometry run of each technical replicate performed twice. The twelve spectral counts for identified proteins were normalised and compared between empty vector control and each NRP1 construct. (A) Initial volcano plots using fold change of each protein over empty vector controls plotted against *p-*values derived from Student’s t-test. NRP1 and co-receptors VEGFR2 and Class A and D plexins are labelled. Underlying data in Supplemental Table 2 (B) Table showing the number of identified proteins in each condition from i) the initial dataset ii) after excluding proteins that were not enriched over empty vector control and iii) which reached statistical significance when *p*-values were assessed using multiple lognormal t-tests adjusted using the two-stage Benjamini, Krieger, & Yekutieli FDR procedure method with an FDR cut-off of 1%. (C) Venn diagram showing the number unique and shared significantly enriched proteins along the two axes of comparison: between each construct in a single biotinylation condition; and between different biotinylation conditions for each single construct.

**Supplemental Figure 3:**
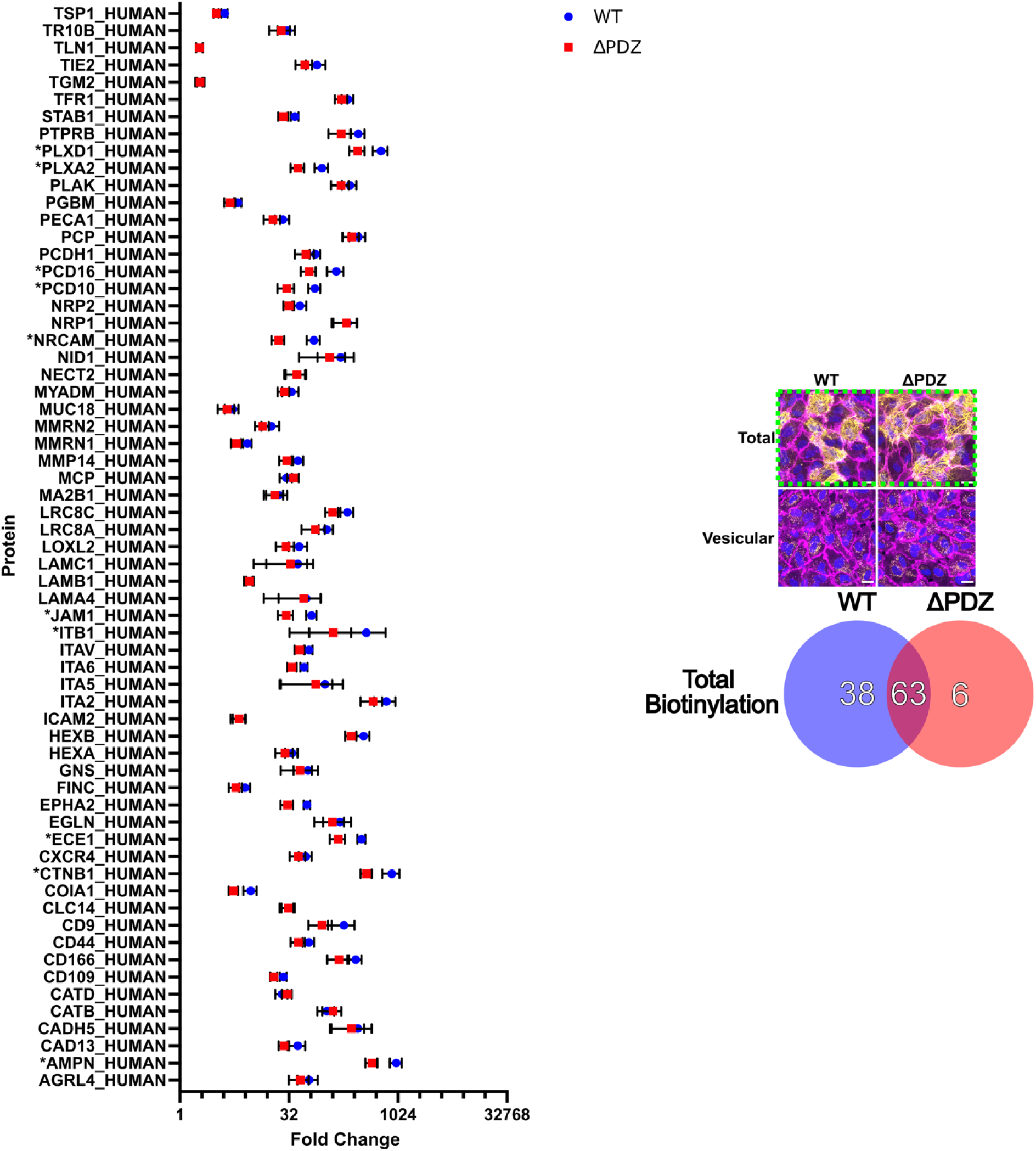
Proteins enriched in both NRP1 WT and ΔPDZ HRP total biotinylation proteomic datasets. HRP proximity biotinylation and mass spectrometry was performed as described above. Proteins with a greater than 2 - fold difference between groups are indicated with a (*). Underlying data in Supplemental Table 3.

**Supplemental Figure 4:**
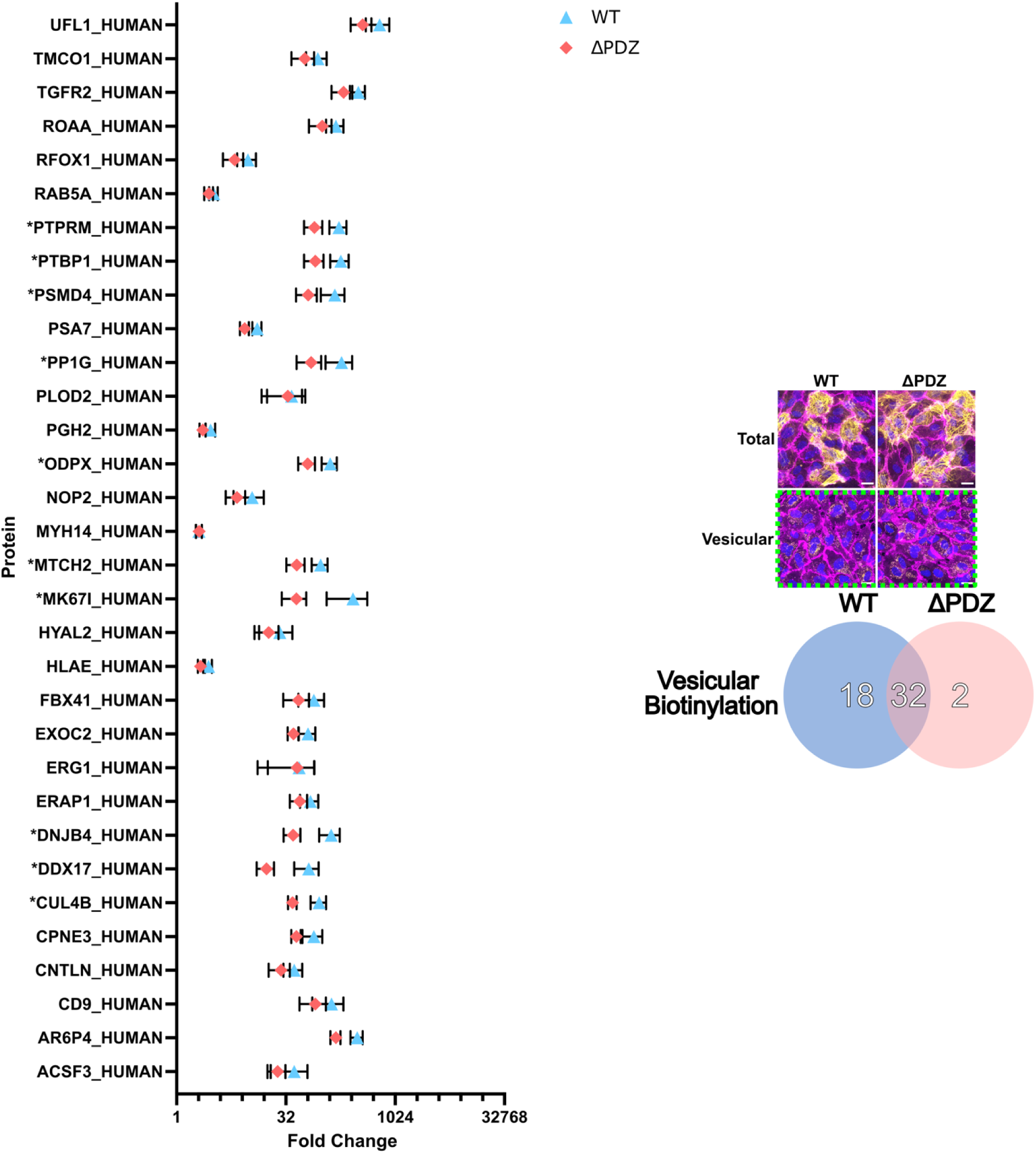
Proteins enriched in both NRP1 WT and ΔPDZ HRP vesicular biotinylation proteomic datasets. HRP proximity biotinylation and mass spectrometry was performed as described above. Proteins with a greater than 2 - fold difference between groups are indicated with a (*). Underlying data in Supplemental Table 4.

**Supplemental Figure 5:**
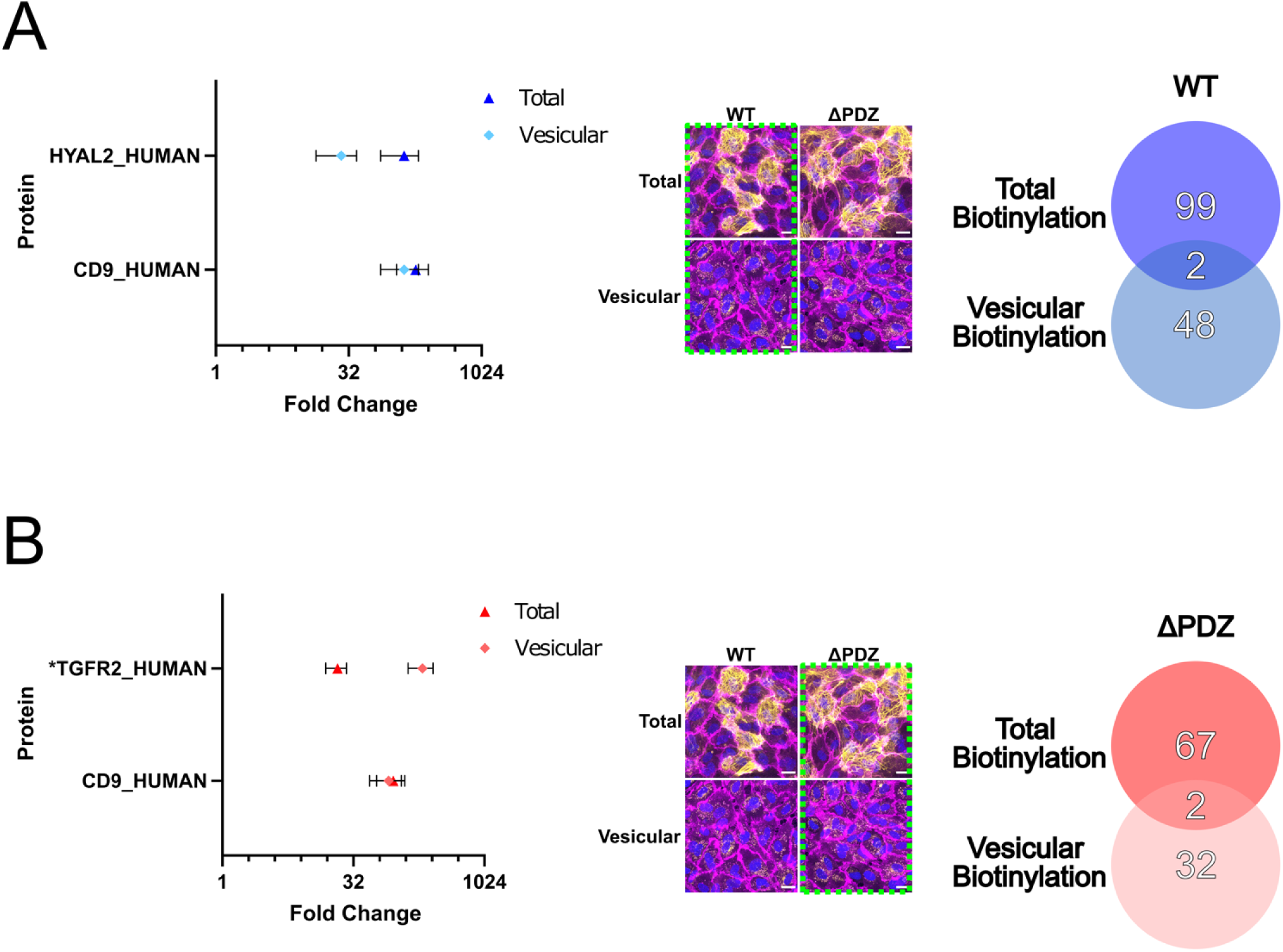
Proteins enriched in both total and vesicular biotinylation proteomic datasets for either NRP1 WT and ΔPDZ HRP. HRP proximity biotinylation and mass spectrometry was performed as described above. (A) Proteins enriched in both total and vesicular biotinylation proteomic datasets for NRP1 WT HRP. (B) Proteins enriched in both total and vesicular biotinylation proteomic datasets for NRP1 ΔPDZ HRP. Proteins with a greater than 2-fold difference between groups are indicated with a (*). Underlying data in Supplemental Tables 5 and 6.

**Supplemental Table 1:** Antibodies used in this study. *Anti-sheep antibody used as secondary antibody against goat primaries

| Antibody | Supplier | Supplier ID | Application |
|---|---|---|---|
| rabbit anti-NRP1 | Abcam | ab81321 | Immunofluorescence |
| rabbit anti-GFP | Santa Cruz Biotechnology | Sc-8334 | Immunofluorescence |
| mouse anti-VE-Cadherin | Santa Cruz Biotechnology | Sc-9989 | Immunofluorescence |
| Streptavidin Alexa Fluor 647 | Thermo Fisher Scientific | S21374 | Immunofluorescence |
| Goat anti-PECAM-1 | Thermo Fisher Scientific | Sc-1506 | Immunofluorescence |
| Donkey anti-rabbit Alexa Fluor 488 | Thermo Fisher Scientific | A21206 | Immunofluorescence |
| donkey anti-mouse Alexa Fluor 555 | Thermo Fisher Scientific | A31570 | Immunofluorescence |
| donkey anti-sheep Alexa Fluor 555* | Thermo Fisher Scientific | A21436 | Immunofluorescence |
| rabbit anti-NRP1 | Cell Signalling Technology | 3725S | Western Blot |
| rabbit anti-VEGFR2 | Cell Signalling Technology | 9698 | Western Blot |
| mouse anti-β-tubulin 2.1 | Sigma Aldrich | T4026 | Western Blot |
| rabbit anti-GAPDH | Sigma-Aldrich | G9545 | Western Blot |
| Streptavidin poly-HRP | Thermo Fisher Scientific | 21140 | Western Blot |
| goat anti-mouse IRDye 680RD | LICORbio | 926-68070 | Western Blot |
| goat anti-rabbit HRP | Agilent | P0448 | Western Blot |

